# Genome-wide analyses of histone modifications and chromatin accessibility reveal the distinct genomic compartments in the Irish potato famine pathogen *Phytophthora infestans*

**DOI:** 10.1101/2022.02.18.480484

**Authors:** Han Chen, Haidong Shu, Yufeng Fang, Wenrui Song, Zhi Li, Yujie Fang, Yuanchao Wang, Suomeng Dong

## Abstract

*Phytophthora infestans*, the causal agent of potato late blight, is a devastating plant disease that leads to Irish potato famine and threatens world-wide food security. Despite the genome of *P. infestans* has provided fundamental resource for studying the aggressiveness of this pandemic pathogen, the epigenomes remain poorly understood. Here, utilizing liquid chromatography-tandem mass spectrometry (LC-MS/MS), we demonstrate post-translational modifications (PTM) at *P. infestans* core histone H3. The PTMs not only include these prevalent modifications in eukaryotes, and also some novel marks, such as H3K53me2 and H3K122me3. We focused on the trimethylations of H3K4, H3K9 and H3K27 and H3K36, and profiled *P. infestans* epigenomes employing Native Chromatin Immunoprecipitation followed by sequencing (N-ChIP-seq). In parallel, we mapped *P. infestans* chromomatin accessibility by Assay for Transposase-Accessible Chromatin with high-throughput sequencing (ATAC-seq). We found that adaptive genomic compartments display significantly higher levels of H3K9me3 and H3K27me3, and are generally in condense chromatin. Interestingly, we observed that genes encoding virulence factors, such as effectors, are enriched in open chromatin regions that barely have the four histone modifications. With a combination of genomic, epigenomic, transcriptomic strategies, our study illustrates the epigenetic states in *P. infestans*, which will help to study genomic functions and regulations in this pathogen.

**Author summary:** Epigenetics play an important role in various biological processes of eukaryotes, including pathogenicity of plant pathogens. However, the epigenetic landscapes are marginally known in oomycetes that are fungal-like organisms and comprise lots of destructive plant pathogens. In this study, using the Irish potato famine pathogen *Phytophthora infestans* as a model, we conducted genome-wide studies of histone post modifications and chromatin accessibility, and demonstrate the relationship of gene expression and evolution with the epigenetic marks. We found that one of the most important classes of virulence proteins, effectors, are enriched in open chromatin regions that barely have eu- and hetero-chromatic marks. This study provides an overview of the oomycete epigenetic atlas, and advances our understanding of the regulation of virulence factors in plant pathogens.

## Introduction

Late blight, a devastating disease is caused by *Phytophthora infestans* that strikes tomatoes and potatoes. The disease is notorious for trigging the 1840s Irish potato famine that resulted in the death of roughly one million people and the displacement of another million [1–3]. Today, *P. infestans* still remains a big threat to global plant health, causing ∼US$6.5 billion losses annually [4–6]. Physiologically and morphologically *P. infestans* resembles filamentous fungi; however, it belongs to oomycete in the Stramenopila kingdom [7, 8]. A large number of oomycetes are destructive plant pathogens whose virulence extensively reply on virulence factors including extracellular toxins, hydrolytic enzymes and inhibitors, and effectors that can enter the cytoplasm of plant cells [9]. In particular, effectors are one of the most important factors that alter host physiology via initiating and allowing an infection to develop [10].

*P. infestans* has a large and complex genomes (∼240 Mb), approximately 74% of the genome is composed of repeats, such as transposable elements [11]. Due to fast bursts of the transposable element activities, repeat-rich genomic regions are highly dynamic and prone to evolutionary changes at accelerated rates compared with the rest of the genome, which shapes the *P. infestans* genome “two-speed” [9, 12, 13]. Interestingly, these repeat-rich regions tend to harbor genes that contribute to virulence such as effector genes, modification enzymes were identified [9, 12, 13]. To date, studies have shown functions and evolutionary trajectory of the effector genes (reviewed in [9, 12, 14]), but how these genes are regulated is marginally known. Epigenetic regulation is crucial for gene expression in the eukaryotes, which is generally influenced by chromatin states, non-coding RNA (ncRNA) and associated machinery, and modifications of DNA or histones, etc. [15, 16]. Importantly, pathogenicity traits, such as virulence [17], sexual reproduction [18], growth [19, 20], and drug resistance [21] have been shown to correlate with epigenetic regulation in eukaryotic pathogens. For instance, in the fungal kingdom, histone modifications, ncRNA mediated silencing and chromosome alterations have been shown to regulate the expression of virulence factors, antifungal drug resistance, and interaction with hosts during infections [21–26]. Similarly, in parasites, histone methylations were reported to control growth and virulence [17, 27]. In oomycetes, epigenetic studies are limited, but studies indicated that epigenetic profiles of oomycetes are distinctive comparing to other organisms, and that ncRNA, histone modifications and chromosome states contributed to virulence. Strikingly, the prevalent 5-cytosine methylation (5mC) is missing in *Phytophthora* species, instead 6-adenine methylation (6mA) is widely distributed across the genomes [28]. This unusual DNA methylation profile makes *Phytophthora* a great model to study the evolution of DNA methylation. Recently, ChIP-seq based on the H3 variant CenH3 (CENP-A) uncovered the centromeres in *P. sojae*, which lack H3K4me2 but embed within the heterochromatin marks H3K9me3 and H3K27me3 [29]. *In silico*, several histone modification enzymes were identified in *Phytophthora*, including histone acetyltransferases (HATs), deacetylases (HDACs) [30], some of which were involved in metabolic and biosynthetic process, sexual reproduction and virulence [31, 32]. In addition, gene silencing has been shown to be mediated by multiple different mechanisms, and be an effective way for oomycete pathogens to modulate virulence factors [33–35]. In *P. sojae,* transcription of the effector *Avr3a* was regulated by sRNA, which led to transgenerational silencing [33], while silencing of another effector gene *Avr1b* was correlated with H3K27me3 deposition [36]. In comparison, transgene-induced silencing of the elicitor gene *INF1* involves chromatin alteration in *P. infestans*, *INF1*- silenced strains harbor distinctive chromatin accessibility in the *INF1* loci [37, 38].

Despite evidence indicated that epigenetic processes affect growth, reproduction, and virulence, we lack an overview of the epigenetic states especially the histone modifications and chromosome states in oomycetes. In this study, we used *P. infestans* as a model to systemically investigate the modifications of histone H3 by LC-MS/MS. We focused on four important H3 modifications, H3K4me3, H3K9me3, H3K27me3 and H3K36me3 that are hallmarks of eu-and hetero- chromatins, and profiled their distributions across the genome. We also assessed the chromatin accessibility via ATAC-seq. We demonstrate that the histone H3 methylations and chromatin accessibility reflect *Phytophthora* genome structure, evolution and gene expression, and correlate closely with virulence factors. These findings provide a new insight of oomycete epigenome structures, and advance our understanding of oomycete genome architectures, which sheds the light on future studies aimed at the epigenetic regulation of plant pathogens.

## Results

### Detection of histone H3 post-translational modifications (PTMs) in *P. infestans*

We are interested in dissecting how *P. infestans* histone H3 are decorated. A previous study has cloned *P. sojae* H3 (PsH3), and demonstrated its predominant nuclear localization in the *P. sojae* transformants by fusing to GFP [39]. To identify histone H3 in *infestans*, we performed blast searches against *P. infestans* T30-4 genome employing the PsH3 ortholog. We found five H3 homologs (PITG_03551, PITG_05675, PITG_20725, PITG_06953 and PITG_13828) present in *P. infestans*. Closer examination of each H3 homologs revealed that PITG_13828 is CENP-A (CenH3), because of its sequence and structure similarity to the *P. sojae* CENP-A [29] (data not shown). Intriguingly, PITG_03551 and PITG_05675 (PiH3-1) have an identical amino acid sequence, but different nucleotide sequences, so did for PITG_20725 and PITG_06953 (PiH3-2) (S1 Fig., S2 Fig.).

To detect histone H3 post-translational modifications (PTMs) in *P. infestans*, we conducted high-performance liquid chromatography-mass spectrometry (HPLC-MS) employing trypsin digested histones. In total, we detected 23 PTM forms in *P. infestans* (Fig. 1A). Most reported H3 PTMs can be found over the *P. infestans* H3 tails, such as the conventional mono-, di-, and tri- methylation of lysine [K], acetylation of lysine, tyrosine and serine [K/T/S], and recently discovered butyrylation [K], 2- hydroxyisobutyrylation [K], crotonylation [K], hydroxylation [Y], and malonylation [K]. Of note, we did not detect the acetylation of K27 that was a common acetylation form in other organisms. On the other hand, we found some unique PMTs, such as di- methylation at K53, tri-methylation at K122 (S3 Fig., S4 Fig.).

**Fig. 1.**
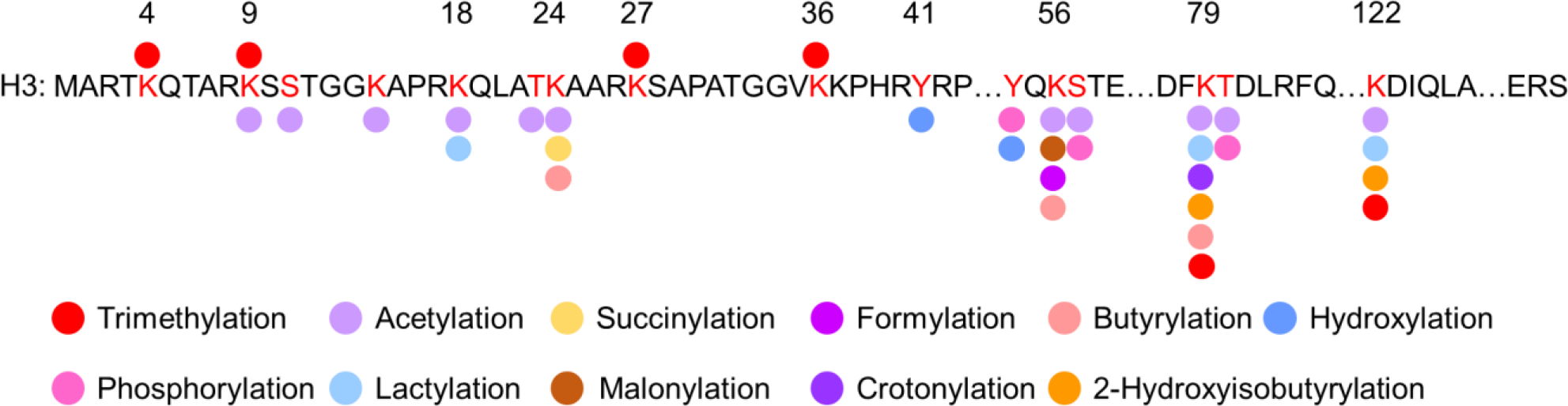
Histone H3 modifications identified in *P. infestans*. Schematic showing various *P. infestans* H3 (PiH3) PTMs that was detected by HPLC-MS/MS. Numbers at top of the PiH3 amino acid sequence indicate the positions of residues. Dots with different colors denote different types of PTMs, and the red ones at the top of K represent lysine tri- methylation.

Further examination revealed distinct distributions of PTMs over the two H3 orthologues. Five PTM variations occurred across 10 amino acid substitutions between two types of H3 (S3 Fig.). Acetylation and phosphorylation at T32 were detected in PiH3-1 Besides, acetylation at T31, di-methylation at K53 and phosphorylation at Y78 were detected in PiH3-2. It suggests different biological functions of H3 variants. Taken together, these findings indicate that despite most of the histone PTMs are conserved in *P. infestans* H3, several distinctive modification patterns are detected over several residues.

### Profiling the genome-wide distribution of four histone H3 methylations and chromatin accessibility

To study the function of H3 PTMs, we focused on four histone methylations H3K4me3, H3K36me3, H3K9me3 and H3K27me3, which are the hallmarks of transcriptionally active euchromatin and transcriptionally silent heterochromatin. To validate the expression of the four histone H3 methylations in *P. infestans*, we carried out western blot using antibodies against these marks. As shown in S5 Fig., all the four histone methylations were detected in *P. infestans*, but not in the *Escherichia coli* expressed H3 protein, as it lacks post histone modifications.

To study genome-wide distribution of H3K4me3, H3K36me3, H3K9me3, and H3K27me3, we performed native chromatin immunoprecipitation (N-ChIP) employing the four antibodies that were tested in western blot, followed by high-throughput Illumina DNA sequencing. We generated high-quality map of each histone methylation with an average of 30 million reads that were uniquely mapped to the *P. infestans* reference genome (S6 Fig.). The four histone methylations displayed distinct distributions: For the euchromatic marks, the majority of H3K4me3 peaks (62.6%) were distributed in gene bodies, while 9.86% were in upstream of genes and 25.25% in intergenic regions (Fig. 2A). Moreover, we found thatH3K4me3 was highly enriched in transcription start sites (TSS) (Fig. 2B). In comparison, H3K36me3 signals were scarce in upstream regions, but were highly enriched in gene bodies (41.46%) and intergenic regions (54.02%) (Fig. 2A and B). With respect to the heterochromatic marks H3K9me3 and H3K27me3, they generally exhibited a similar distribution pattern: 95.44% H3K9me3 peaks and 96.01% H3K27me3 peaks were localized in the intergenic regions (Fig. 2A, B and C).

**Fig. 2.**
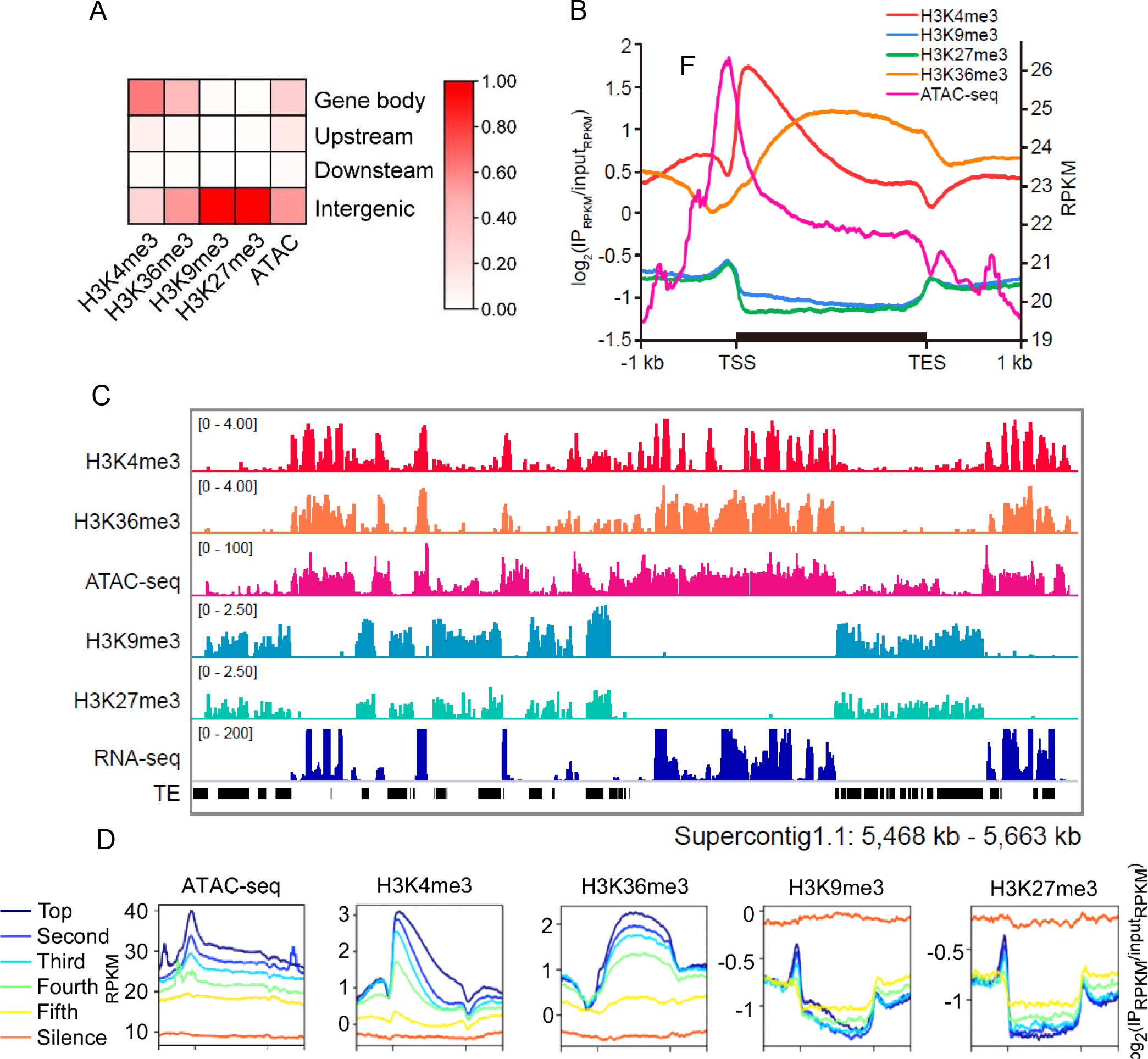
Distribution of four histone H3 methylations and chromatin accessibility. (A). Heatmap showed the distribution of ChIP-seq and ATAC-seq peak. upstream was defined as -800 bp before start code, downstream was defined as +400 bp after stop code. Submits of peaks in S6C Fig. were used. (B). line plot showed the distribution of four histone H3 methylations and chromatin accessibility across gene, -1 kb to +1 kb region were included. (C). Representative snapshot of H3 methylations and ATAC-seq statue. The snapshot was collected from IGV software. (D), Accessible chromatin, H3K4me3 and H3K36me3 are associated with highly expressed genes, meanwhile, H3K9me3 and H3K27me3 are associated silenced genes.

One remarkable feature of heterochromatic marks is its association with repetitive sequences [40]. In *P. sojae*, repetitive sequences, in particular, transposable elements (TEs) were highly enriched with H3K27me3 and H3K9me3 [41]. Similarly, we found that TEs in *P. infestans*, were highly enriched with H3K9me3 and H3K27me3, while lacked H3K4me3 and H3K36me3 (Fig. 2C and S7A Fig.). Further investigation of different TE families revealed that 55% H3K9me3 and 66% H3K27me3 signals were associated with LTR retrotransposons that is the major TE family is the *P. infestans* genome (S7B Fig.). Interestingly, the two heterochromatic marks exhibited quite different distribution over the DNA transposons, with 16% H3K9me3 and 5% H3K27me3 (S7B, Fig).

To further investigate the chromatin state over the *P. infestans* genome, we profiled the chromatin accessibility based on Assay for Transposase-Accessible Chromatin with high-throughput sequencing (ATAC-seq). A high-quality chromatin accessibility map was generated by aligning 100 million ATAC-seq reads uniquely to the reference genome (S6 Fig). We found that almost half of the ATAC-seq reads were mapped to the intergenic regions, while 29.05% were in gene bodies and 12.59% were in upstream of genes (Fig. 2A). To examine the relationship between chromatin accessibility and the four histone modifications, we implemented correlation analysis by PCA and heatmap. We found that heterochromatin was anticorrelated with chromatin accessibility (Pearson co-efficiency ranging from -0.15 to -0.23), while half euchromatin were somewhat correlated with chromatin accessibility (Pearson co-efficiency ranging from 0.12 to 0.20) (S8 Fig, S9 Fig). Comparing to H3K4me3, we found that the major ATAC-seq signals approximal gene bodies were around 100 bp upstream of TSS, indicating that the open chromatin state of promoter regions (Fig. 2B). It suggests the unique role of chromatin accessibility in chromatin state defining compared with those four PTMs.

### Chromatin states strongly correlate with gene expression

We next sought to examine the correlation of H3 modifications and chromatin accessibility with gene expression. To determine expression levels of genes from the same growth stage that was used for profiling chromatin states, we performed RNA-seq from the *P. infestans* mycelial stage. We classified all *P. infestans* genes into six categories from “Silent” to “Top” expression, based on their expression levels, (Fig. 2D). We found that genes of higher expression were generally located in chromatin regions that are more accessible, and that H3K4me3 and H3K36me3 were positively correlated with gene expression (Fig. 2D) suggesting open chromatins and these euchromatic histone marks are associated with active genes. In contrast, H3K9 and H3K27 tri- methylations exhibited an anticorrelation with expression level, indicating these two histone marks are associated with silencing (Fig. 2D). Genome browser views of individual genomic locations also suggest a strong association of transcription activity with chromatin accessibility and the four histone marks (Fig. 2C).

Collectively, these results illustrate that methylation of H3K4 and H3K36 is associated with expressed genes, which are correlated with accessible chromatins; while the methylation of H3K9 and H3K27 is enriched in areas of the genome that are transcriptionally silent and are generally consisted of TEs, which tend to be closed chromatin conformation.

### Chromatin states reflect the bipartite genome structure

The genomes of *Phytophthora* species show a bipartite “two-speed” architecture that is consisted of gene-dense regions (GDRs), and gene-spare regions (GSRs) [11, 13, 42]. The GSR compartments are associated with accelerated gene evolution, serving as a cradle for adaptive evolution [11, 13, 42]. To investigate the relationship between chromatin state and the genome architecture, we measured the average level of each epigenetic state for genes located in GDRs and GSRs. We discovered that GSRs were enriched with the heterochromatic marks H3K9me3 and H3K27me3, and had condense chromatins, but lacked the euchromatic marks H3K4me3 and H3K36me3 (Fig. 3). In contrast, GDRs were enriched with the euchromatic marks H3K4me3 and H3K36me3, and were associated more accessible chromatin (Fig. 3). Thus, the epigenetic states display close correlation with the “two-speed” genome architecture.

**Fig. 3.**
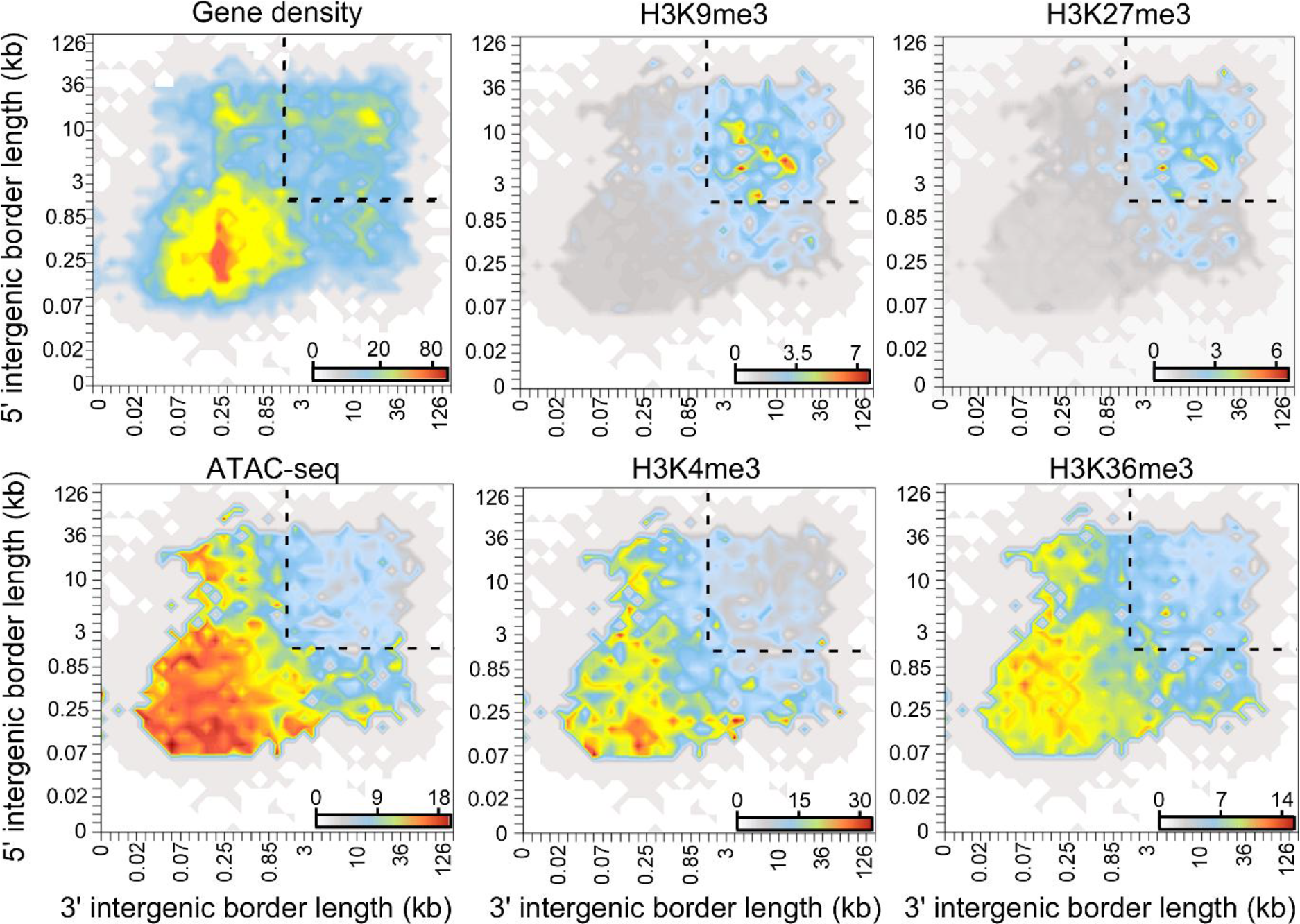
Distribution of H3 methylations and chromatin accessibility reflects the bipartite genome structure. ATAC-seq, H3K4me3 and H3K36me3 density in GDR. The dotted line highlights GSR. Gradient color represents gene number is gene density plot, and gradient color in other plot represent the average normalized ATAC-seq or H3 methylation signal.

### Chromatin states are associated with the conservation of protein-coding genes

To address the connection between epigenetic state and protein evolution, we first sought to identify genes that were evolutionarily conserved in *P. infestans*. We selected 17 eukaryotic species including oomycetes, fungi, plants and animals, and measured the ratios of non-synonymous/synonymous codon substitution (dN/dS) for protein-coding genes. We found that the most conserved genes across eukaryotes had lowest dN/dS ratios, while the genes specific to *P. infestans* had highest dN/dS ratios, suggesting species-specific genes are evolved later (S10A, B Fig.). In agreement with this observation, high dN/dS density was also found in GSRs, further implying that the genes in these regions were under higher evolutionary pressure (S10C Fig.).

Based on the gene evolution analysis, we examined the epigenetic states over each protein-coding gene in *P*. *infestans*. We found that *P. infestans* specific genes had higher level of H3K9me3 and H3K27me3, and were generally associated with condense chromatin. In contrast, conserved genes had higher level of H3K4m3 and H3K36me3, and were preferably associated with more open chromatin regions (Fig. 4A). We found that genes that were under positive selection accumulated more H3K9me3 and H3K27me3 signals, whereas the genes under purifying selection accumulated more H3K4me3 and H3K36me3 (Fig. 4B). These findings are consistent with the chromatin accessibility and methylation pattern of the “two speed” genome architecture that we found above, namely the conserved genes are usually distributed in accessible GDR, and thus are marked by H3K4me3 and H3K36me, whereas the rapidly evolved TEs are enriched in the condensed GSR and thus associated with H3K9me3 and H3K27me3.

**Fig. 4.**
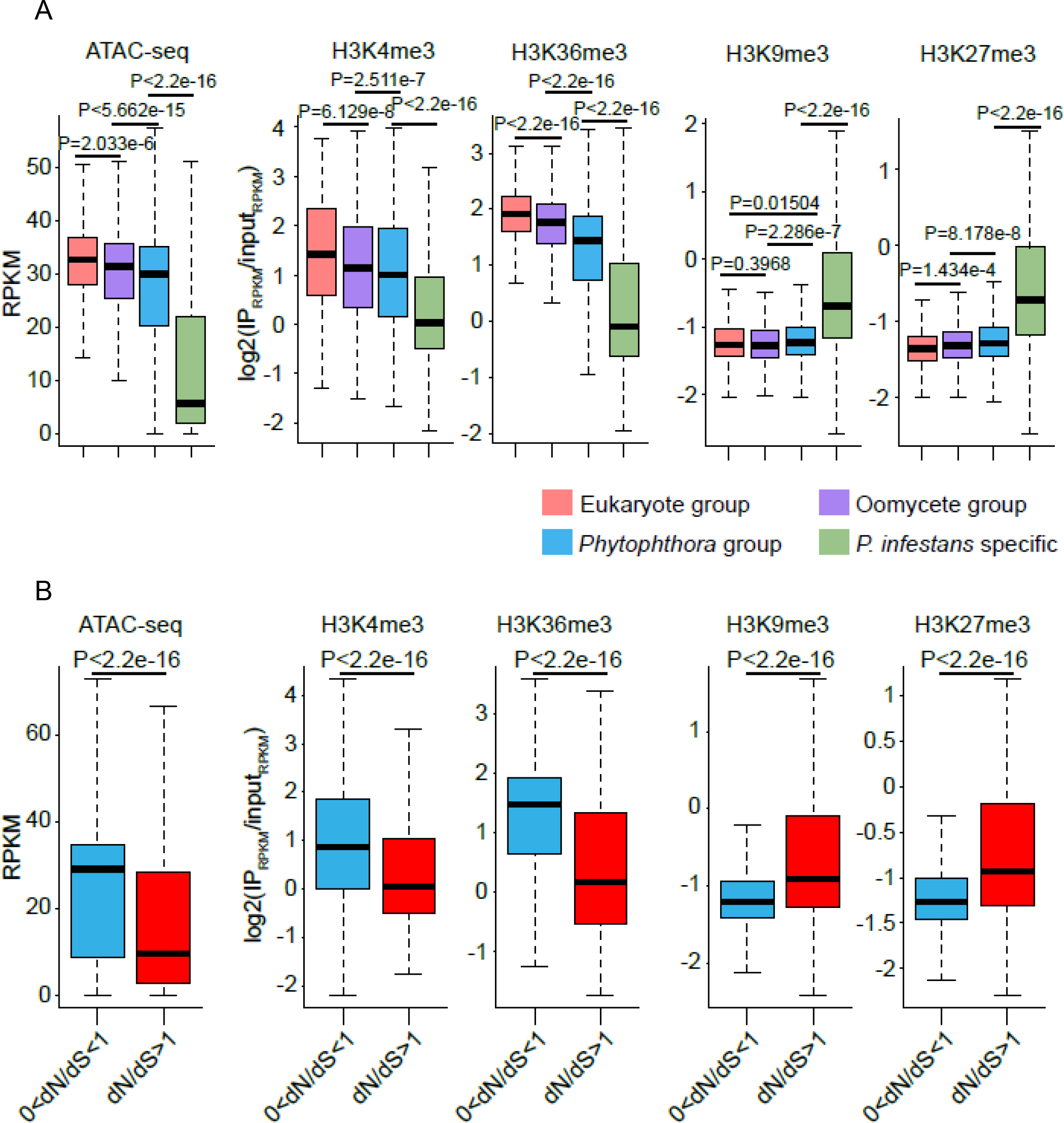
Histone H3 methylations and chromatin accessibility are associated with evolution of the protein-coding genes. H3K9me3 and H3K27me3 were abundant in *P. infestans* specific genes (A) and fast evolved genes (B) with generally condense chromatin, whereas higher H3K4m3 and H3K36me3 preferably associated with conserved genes which were located in more accessible chromatin regions. In (A), Pink, purple, blue and green represent eukaryote, oomycete, Phytophthora, and P. infestans specific gene groups, respectively. In (B), all protein-coding genes were divided into two categories based on their dN/dS ratios, their methylation and ATAC-seq density were compared. The number of genes in two groups were 14291 (0<dN/dS<1) and 1213 (dN/Ds>1), respectively.

### Effector genes are enriched in highly accessible chromatin region with less histone methylation level

To characterize the functions of genes that were undergone histone methylation, we performed gene ontology (GO) analysis. We found that histone methylations widely contributed to biological processes (S11 Fig.). Interestingly, while the processes regulated by the euchromatic marks H3K4me3 and H3K36me3 were mostly different, these regulated by H3K9me3 and H3K27me3 were largely coincident (S11 Fig.), further indicating cross-talks may occur between the two heterochromatic marks for regulating gene expression. Notably, GO analysis indicated that histone modifications broadly contribute to pathogenesis related functions, such as pectate lyase activity and extracellular region, indicating the epigenetic regulation are critical for virulence. This prompted us to investigate the relationship between histone methylations and secreted proteins (secretome), in particular RxLR effectors [43, 44]. We found that comparing to other protein-coding genes, genes encoding secretome and RxLR effectors had overall higher H3K9me3 and H3K27me3and lower H3K4me3 and H3K36me3 densities (S12A, and S12B Fig.). It is possibly because the majority genes encoding RxLR effectors were repressed in mycelia stage.

To further investigate the role of chromatin state in regulating virulence, we generated a multivariate Hidden Markov Model based on the distribution of the four aforementioned histone marks and chromatin accessibility by chromHMM. *P*. *infestans* genome was divided into five distinct states including open chromatin (OC) that harbored ATAC-seq signals but lacked the four histone marks; strong transcription (ST) region that had abundant H3K4me3, H3K36me3, and ATAC-seq signals; H3K9me3 dominant repression region (H3K9DR); H3K27me3 dominant repression region (H3K27DR); and quiescent (Quies) state that were absent of all the tested marks (Fig 5A). We found that 53.7% (868/1616) genes encoding secreted virulence factors, such as carbohydrate-active enzymes (CAZyme) and various effector families such as Crinkler effectors (CRN), necrosis- and ethylene-inducing-like proteins (NLP), small cysteine rich effector proteins (SCR) and RxLR, were associated with the OC and ST states (Fig. 5D, E). Interestingly, despite OC state accounts for the smallest percentage (5.29% of the genome), 44% RxLR effector-coding genes were associated with this state (Fig. 5E). A further analysis showed CAZyme, NLP and RxLR gene families significantly enriched in OC state (Fig. 5F). GO analysis of genes from different chromatin state revealed, the GO items like extracellular region, cell wall organization and pectin catabolic process were the most enriched ones in OC state associated genes (S13 Fig.), suggesting the role of OC state in *P. infestans* pathogenicity gene regulation.

**Fig. 5.**
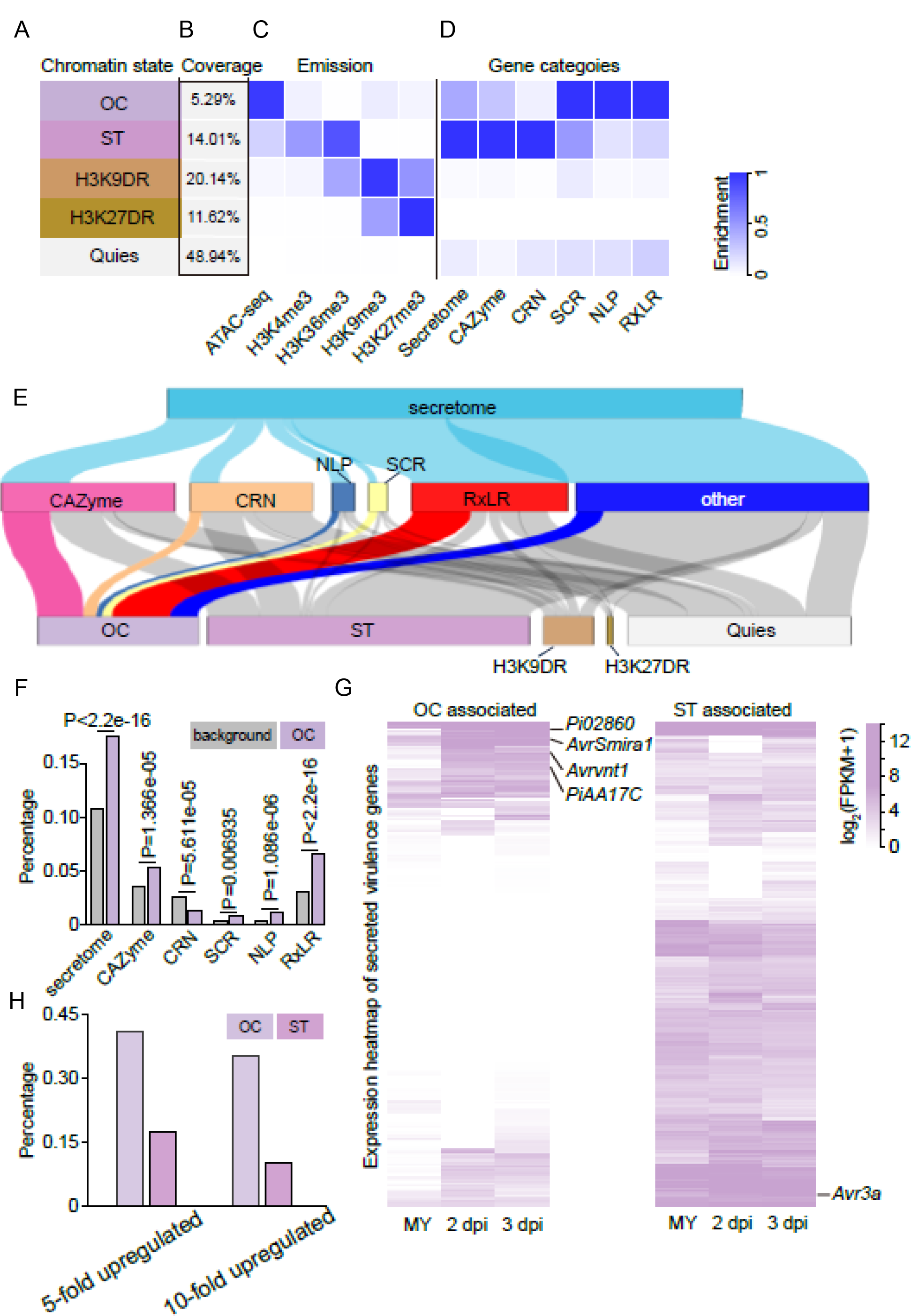
Virulence genes were enriched in OC state. A. Chromatin state definitions, open chromatin (OC), strong transcription region (ST), H3K9me3 dominant repression region (H3K9DR), H3K27me3 dominant repression (emission probability) of histone marks and chromatin accessibility were presented. D, Enrichments probability of different gene categories in each chromatin states measured by chromHMM. E, Secretome genes were divided into CAZyme, CRN, SCR, NLP, RxLR and other genes, and Sankey plot showed chromatin state distribution of those gene groups. F, Percentage of gene in OC states compared with background group. Grey bars were calculated as (gene number of corresponding gene group)/ (total gene number of *P. infestans*). Purple color bars were calculated as (number of gene in corresponding gene groups associated with OC state)/ (total number of genes associated in OC state). P-value was calculated by Chi-squared test. G, Gene expression heatmap of OC state and ST state associated secreted virulence genes in mycelium (MY), 2-day post incubation (2 dpi) and 3-day post incubation (3 dpi) stages. Expression heatmap were normalized by log_2_(FPKM+1) and clustered in row scale by Tbtools. H, Summary of percentage by gene upregulation fold change in OC state and ST state associated secreted virulence genes, the gene upregulation fold change was calculated using row FPKM value.

To address how the expression profiles of the virulence genes correlated with the OC state. We examined the transcriptomes of the *P. infestans* T30-4 strain at different life/infection stages, such as mycelium (MY), 2 dpi (days post incubation) and 3 dpi. We found that about half of the OC state-associated secreted virulence genes (CAZyme, CRN, NLP, SCR and RxLR) displayed infection-induced pattern, for instance secreted pectin monooxygenase encoding gene *PiAA17C* [45], and RxLR effector genes *Avrvnt1*, *AvrSmira1 and Pi02860* [46–48] which are key virulence genes. In contrast, most ST state-associated secreted virulence genes were relative highly expressed, although some of them were infection-induced like Avr3a (Fig. 5G). To further explain the expression difference, we summarized the upregulation fold change of OC state and ST state-associated secreted virulence genes. Among 251 secreted virulence genes in OC state, 41.03% genes were 5-fold upregulated, and 35.45% genes were 10-fold unregulated at least in one infection stage. Otherwise, the ratio is lower in ST state (Fig. 5H). It suggests that OC state-associated virulence genes could be highly induced in infection stages. Altogether, our analyses implied the importance of OC state in virulence gene regulation.

## Discussion

In this study, we identified the histone H3 PTMs in *P. infestans* by HPLC/MS, and generated an epigenetic atlas on basis of histone H3 methylations and chromatin accessibility. We found that most H3 PTMs were conserved in *P. infestans*. By examining the tri-methylation profiles of H3K4, H3K36, H3K9 and H3K27 and chromatin accessibility, we concluded that these chromatin states were closely associated with gene expression, genome structure, protein evolution, and virulence factors regulation. Our results provide evidence that epigenetic scenario is associated with pathogen genome in oomycete, and highlight chromatin state of virulence factors.

We found *Phytophthora* H3 proteins and H3 PTMs are slightly different from these reported in animal, plants and Protista, indicating dynamics of these highly conserved protein. With blast search, we found two H3 orthologues in *P. infestans*; however, with sequence composition and phylogenetic analyses, we cannot distinguish the canonical orthologue H3.1 and the variant H3.3 in *Phytophthora* (S2 Fig.). On the other hand, HPLC/MS revealed differences of PTM between the two *P. infestans* H3 orthologues (S3 Fig). This suggests that oomycete histone H3 may have a distinct evolutionary trajectory, additional experiment such as gene expression pattern examination in and outside of the S phase, and nucleosome assembly will help to clarify the two H3 variants in *Phytophthora*.

We found some unique modifications present in *P. infestans*, such as methylations at residues H3K53 and H3K122. H3K53me maybe species-specific, as in several organisms the residue 53 in H3 not lysine but arginine (S2B Fig.). In *Arabidopsis*, the H3K53 site was also reported to possess a modification; however, instead of methylation, it was 2-hydroxyisobutyrylation (Khib). Intriguingly, this unusual Khib modification contributed to the plant adaption to stress [49], suggesting H3K53me2 may also participate in some biological activities in *P. infestans*. Despite studies have shown that H3K122ac is a critical transcriptional regulator that defines genome-wide genetic elements and chromatin features associated with active transcription in mammalian cells [50], there is no report of H3K122 methylation, thus it will be of interest to examine its function in oomycetes. While most histone H3 PTMs reported to date can be found *in P. infestans*, we found acetylation of H3K27 and H3K36 are missing in the pathogen. Interestingly, histone acetyltransferase families that represent for H3K27ac and H3K36ac were identified [30], and a recent paper found both H3K27ac and H3K36ac in *P. infestans* [51]. It is possible that this modification level was too low to be detected in our mass spectrometry. Overall, we confirmed that large number of PTMs were presented in *P. infestans*, meanwhile, some unique PTMs were detected.

In *P. sojae*, it has been shown that H3K9me3 and H3K27me3 are generally coincident over the genome [29]. We generally observe a similar distribution pattern in *P. infestans*. Surprisingly, closer examination of TE demonstrated that the two heterochromatic marks were separated over the DNA transposons (S7 Fig.), implying a different mode may be used for silencing the types of TE. Another interesting features of epigenetic marks we found in *P. infestans* is that H3K36me3 distributed over the gene bodies of active genes (Fig. 2B), which is distinctive from strong methylation peak at the 3’ end of the gene body in human and mouse, and strong peak at the 5’ end of the gene body *Arabidopsis* and rice, but is similar to the pattern in the brown alga *Ectocarpus siliculosus*, yeast, and *C. elegans* [52, 53], suggesting divergent regulators and mechanisms for establishing H3K36 methylation among species.

Genome-wide profiling of chromatin states provides insights into the unique genomic compartments of plant pathogens. Principally, here we directly connected the chromatin accessibility, histone methylation status and two-speed genome feature, and greatly enriched the dimension of two-speed genome (Fig. 3). Similar to our results, high enrichment of the heterochromatic marks H3K9me3 and H3K27me3, rather than the euchromatic mark H3K4me2 was found in the rapidly evolved accessory chromosomes in the wheat fungal pathogen *Zymoseptoria tritici* [54]. Recent research in the verticillium wilt fungus *Verticillium dahliae* revealed adaptive lineage-specific genomic regions contain many heterochromatin features, but more accessible than true heterochromatin [55]. The heterochromatin features of adaptive genomic compartments in plant pathogens remains a question that the formation and maintenance of these regions. Mechanically, experimental evolution evidence proved that loss of H3K27me3 by knockout methyltransferases Kmt6 drastically reduces the loss of accessory chromosomes [56]. It explained why these regions are more susceptible to genetic diversity. Further experimental research in *P. infestans* is needed to understand the unique genomic compartments regulation. Besides, open chromatin sites reflect the potential of recombination [57, 58], and *A. thaliana* showed that mutational rates are significantly predicted by some epigenetic modifications [59]. It suggests that epigenetic status is associated with evolutional hotspots. So, it’ s reasonable to infer that chromatin state details directly reflect sequence mutation and epimutation ratio in specific strains. Higher resolution of two-speed genome and the molecular basis of genome architectural reshaping are fascinating points to us.

Dissecting the chromatin state of genes encoding secreted virulence protein is one of the key steps to illuminate *P. infestans* pathogenicity regulation. Interestingly, the GO analysis indicated that gene related to extracellular region and pectin catabolic process are enriched in the OC state (Fig. 5, S13 Fig.), meanwhile, those items was also enriched in H3K9me3 and H3K27me3 methylated genes (S11 Fig.). We inferred that those virulence genes involved in these GO items were not overlap and in different regulation model. Firstly, most virulence genes in OC state had the potential to be induced in infection stages, this kind of genes performed strong induction pattern due to transcriptional releasing in infection stages (Fig. 5G, 5H). Secondly, the other virulence genes were strongly repressed by methylation of histone lysine residues. Those kinds of genes might be epigenetically regulated in Phytophthora [36], which was also observed in other filamentous pathogens [60, 61]. Epigenetic silencing of TE by heterochromatin marks to prevent TE proliferation in the TE-rich region is quiet common in filamentous pathogen genomes [62], and we indeed found adjacent TEs around *RxLR* gene cluster (S12 Fig.). Moreover, it has been accepted that epigenetic marks are tissue- and stage-specific [63–65], revealing epigenomes of different stages and strains can dissect holistic virulence gene regulation model. Here, we propose that functional regulatory elements that contributed to establish and maintain normal chromatin state could be drug targets to defect diseases. In conclusion, our findings reported herein provide clues for virulence gene regulation mechanism investigation and pest management.

## Materials and methods

### Oomycete and fungal cultures

*P. infestans* strain T30-4, and *P. sojae* strain P6497 were used in the study. *P. infestans* was routinely maintained on RSA/V8 medium at 18 ℃ in dark, and *P. sojae* on 10% V8 medium at 25℃ in dark [28].. *M. oryzae* strain Guy11 was regularly cultured in CM medium at 28℃ [66]. LC-MS/MS analysis Core histones were extracted from *P. infestans* mycelia employing EpiQuik Total Histone Extraction Kit (EpiGentek, OP-0006-100), according to the manufacture instruction. After being separated by 12% SDS-PAGE, proteins with ∼ 17 kDa size were extracted and analyzed by mass spectrum at Thermo Fisher. Given that H3 proteins have multiple trypsin digestion site (lysine and arginine), a portion of sample was treated with propionic anhydride (PA) to block trypsin digestion at the lysine sites to get maximal sequencing read coverage. The trypsin digested peptides (derivatized and non-derivatized) were injected to HLC/MS/MS (C18 HPLC column and Q Exactive HF-X Mass Spectrometer), and the acquired MS/MS data was analyzed by the pFind3.0 [67] with default parameters.

### Protein extraction and western blot

To extract oomycete and fungal proteins, mycelia collected from liquid cultures of six days old *P. infestans*, three days old *P. sojae*, and three days old *M. oryzae* were dried by filter paper, and ground with mortars and pestles in liquid nitrogen. 800 µl lysis buffer (1% SDS in TE buffer) was added into every 100 mg pulverized oomycete and fungal tissue. Lysates were mixed by vortex for 30 min at 4℃ 200 µl supernatants were collected after spun prepared for western blot. Protein samples were mixed with protein loading buffer (Beyotime, P0015) and denatured at 95℃ for 10 min. To express PsH3 in *E. coli*, the target gene (Ps_322070) was PCR amplified from *P. sojae*, and was cloned into the plasmid pET32a using Vazyme ClonExpress II One Step Cloning Kit C112. The resulting plasmid pET32a-PsH3 was introduced in *E. coli* BL21. The protein was expressed at 30℃ for 8 hours in LB medium with 0.5 mM isopropyl β-D-Thiogalactoside β (IPTG) according to the manufacture instruction [68]. After sonication and centrifugation, about 100 µl supernatant containing *E. coli* crude protein extract was used for western blot. Antibodies H3K36me3 (abcam, ab9050), H3K27me3 (Millipore, 07-449), H3 (abcam, ab1791), H3K4me3 (abcam, ab8580) and H3K9me3 (abcam, ab8898) were used as primary antibodies against the relevant histone H3 modifications, and goat-anti-rabbit IRDye 800CW antibody (Odyssey, no. 926-32211, Li-Cor) was as asecondary antibody. The signals were exposure by laser imaging system (Odyssey, LI-COR company).

### Phylogenetic analysis

H3 protein sequences from human (HsH3.1, accession number: NP_003520.1, HsH3.2, NP_066403.2, HsH3.3/NP_002098.1, HsCenP-A/NP_001035891.1), *Arabidopsis thaliana* (AtH3.1/NP_563838.1, AtH3.3/ NP_001329167.1, AtCenH3/NP_009564.1), *Neurospora crassa* (NcH3/CAA25761.1) and Saccharomyces cerevisiae (ScH3/NP_009564.1) were downloaded from NCBI. *Phytophthora* H2A sequence were from published paper [69]. Homologous gene blast was obtained by Seqhunter2 with an E value of 10^-5^. Sequence alignment was performed by Bioedit software [70], and phylogenetic analysis was conducted by MEGA5 using neighbor- joining model and 10000 bootstrap replicates [71].

### Native ChIP-seq, ATAC-seq and RNA-seq

Native ChIP experiments were performed as previously described [29, 36, 72]. Input and immunoprecipitated DNA samples were sequenced by BGI company as 50SE. To prepare ATAC-seq, *P. infestans* protoplasts were isolated according to refrenece [73].

Protoplast was stored at -80℃ using Nalgene 5100-0001C, and then sent to BGI for treatment [74]. In brief, lysing the cells and keep the cells on the ice all the time. Add Tn5 transposase to the cell suspension after cell lysis and then purify it. The DNA fragments were sequenced in BGI company as PE150. Infection samples (2 dpi and 3 dpi) were prepared as described [75]. RNA were extracted using Omega Total RNA Kit I according to the manufacturer’s manual, RNA-seq libraries were prepared by BGI company and sequenced by BGISEQ-500.

### High-throughput sequencing data analysis

Both ChIP-seq and ATAC-seq reads were polished by BGI prior to be released, and thus were mapped to the *P. infestans* reference genome directly using Bowtie2 (v2.3.5.1) [76] (see S6 Fig. for mappability). The aligned bam files were sorted and indexed by samtools (version: 1.7) [77]. The ChIP-ed and input samples were analyzed with DeepTools (v3.4.3) [78] ‘‘bamCompare” to calculate normalized ChIP signals (log2[ChIP_RPKM_/Input_RPKM_]),. ChIP-seq data was visualized using TBtools [79] and Integrative Genome Viewer (IGV, v2.8.0) (https://software.broadinstitute.org/software/igv/) [80]. ChIP-seq peaks were defined by MACS2 employing a default model for H3K4me3 , and a “--broad” model for H3K9me3, H3K27me3, and H3K36me3 [81, 82]. Peak overlaps were conducted by “intersectBed” in bedtools [83]. To visualize the ATAC-seq reads, BAM files were converted to bigwig format using “bamCompare” in DeepTools with RPKM normalization. ATAC-seq data was visualized using TBtools [79] and IGV v2.8.0 [80]. The ATAC-seq reads were shifted +4 on the positive strands and -5 on the negative strands using deepTools software, and then ATAC-seq peaks were defined by MACS2 in default model. Peak overlap was conducted by “intersectBed” in bedtools.

RNA-seq reads were polished by BGI prior to be released. To generate mRNA profiles, the RNA-seq reads were aligned to the genomes using HISAT2 (version 2.1.0) [84] , and the resulting files (.bam) were sorted and indexed by samtools (version 1.9) [77]. The bam file was converted to .tdf for visualization using IGV. Gene expression data were calculated by StringTie v2.1.2 [85] and was presented as FPKM values.

### Other analysis

PCA plot and heatmap clustering of ChIP-seq and ATAC-seq was using “plotPCA” and “plotCorrelation” in deepTools. Overlap of ATAC peak and ChIP-seq were conducted by “intersectBed” in bedtools. The Pearson correlation computation was used in heatmap clustering. The chromatin state was defined by chromHMM [86] in “BinarizeBed” model using MACS2 peak files. GO enrichment was analyzed by R package clusterprofile v3.14.3 by universal enrichment analyzer “enricher” and “compareCluster” [87]. Gene conservation analysis among species was performed by OrthoMCL v2.0.9 by default parameters [88]. Protein conservation was based on 17 eukaryotic species including oomycetes, fungi, plants and animals, and divided the *P. infestans* coding-genes into four groups (Eukaryote, Oomycete, *Phytophthora* and *P. infestans*), each of which has 1157, 3064, 5680 and 2483 genes. Eukaryote group contains protein-coding genes that conserved among all 17 species. Oomycete group contains protein-coding genes that conserved among 10 oomycetes besides genes in eukaryote group. Phytophthora group and P. infestans specific groups can be inferred like this. The Sankey plot was performed by R package networkD3 [89].

## Acknowledgments

We are grateful to the Bioinformatics Center at Nanjing Agricultural University for support of the bioinformatics analysis. We thank the NJAU oomycete research group for valuable discussion.

---

S1 Fig. Sequences analysis of *P. infestans* H3 homologs.

(A), Nucleotide sequences alignment of four *P. infestans* H3 homologs. (B), Amino acid sequence alignment of four *P. infestans* H3 homologs.

S2 Fig. Phylogenetic and sequence analysis showed conserved Phytophthora H3 homologs.

(A), Phylogenetic analysis of Phytophthora H3 homologs. H3 homologs from human (HsH3.1/accession number: NP_003520.1, HsH3.2/ NP_066403.2, HsH3.3/NP_002098.1, HsCenP-A/NP_001035891.1), *Arabidopsis thaliana* (AtH3.1/NP_563838.1, AtH3.3/ NP_001329167.1, AtCenH3/NP_009564.1) and budding yeast(ScH3/NP_009564.1) were included in the study., H2A was set as an outgroup. (B), Alignment of H3 variants from *P. infestans* (PITG_05675 and PITG_03551) and *P. sojae* H3 (Ps_284752, Ps_322070 and Ps_476994) with HsH3.1, HsH3.2, AtH3.1, ScH3 and NcH3.

S3 Fig. PTM variants in Phytophthora H3.

(A), amino acid substitutions of two H3 forms were compared and different PTMs were marked. The numbers indicated amino acid position, amino acids at each site were listed and different colors indicated PTMs. (B), Individual mass spectrums of five PTM variants were shown.

S4 Fig. Summary of H3 lysine acetylation and methylation among seven organisms.

Highly conserved lysine were listed, red color indicated this PTM was detected or reported, blue color indicated this PTM was not detected or reported. We used grey color to mark the residue at 53 in human, for this site mutated to R.

S5 Fig. H3K4me3, H3K9m3, H3K27me3 and H3K36me3 were detected in *P. infestans*.

(A), MS spectra of H3K4me3, H3K9m3, H3K27me3 and H3K36me3 in *P. infestans*. (B), H3, H3K4me3, H3K9m3, H3K27me3 and H3K36me3 were detected by WB. *P. infestans*,

*P. sojae* and *M. oryzae* total protein were detected by five different antibodies, and prokaryotic expressed H3 protein of *P. sojae* is the negative control that cannot be modified.

S6 Fig. The good quality and correlation between biological replicates of different histone modifications and ATAC-seq in this study.

(A). The overall analysis of ChIP-seq data and ATAC-seq data. (B). Highly repeatability of two replicates. The genome was divided into 2kb bins and RPKM values of each bin were used to calculate Pearson correlation coefficients. (C). Highly overlapped peak of two replicates.

S7 Fig. TE region have higher H3K9me3 and H3K27me3.

(A). Methylation level of gene and TE were compared. Methylation level was calculated as log2(IPRPKM/inputRPKM). P values are calculated with the two-sample Kolmogorov-Smirnov test. (B). H3K9me3 and H3K27me3 are preferentially associated with LTR and non-LTR type transposon in *P. infestans*. Intersected percentage of peaks length and TE length were calculated.

S8 Fig. PCA and heatmap clustering revealed overall distinct features between ATAC-seq and ChIP-seq reads.

PCA analysis (A) and heatmap clustering (B) of bw file generated by deepTools. each dot represents one replicates. The Pearson correlation computation was used in (B).

S9 Fig. Unique role of chromatin accessibility in chromatin state defining compared with those four PTMs by peak analysis.

ATAC peaks were divided into ATAC peak-alone, ATAC and H3K4me3 peak cobound, and etc. The number of peaks were 10666, 3334, 1311, 2496, 1496, 331, 486 and 663, respectively. Overlapped peaks defined in S6C Fig. were used here.

S10 Fig. Genes under positive selection have higher H3K27me3 and H3K9me3 in P. infestans.

(A), Conserved genes have lower dN/dS value, and *P. infestans* specific gene is under highest positive selection. P values are calculated with the two-sample Kolmogorov- Smirnov test. (B). GSR has higher dN/dS ratio. Gradient color represents dN/dS ratio. (C), Genes under positive selection have higher H3K9me3 and H3K27me3. RPKM value from -1 kb to 1 kb region of each gene were collected and the methylation level was calculated as log2(IP_RPKM_/input_RPKM_). P values are calculated with the two-sample Kolmogorov-Smirnov test.

S11 Fig. Comparison of the GO enrichment result of methylated genes.

S12 Fig. Genes encoding secretome protein and RxLR effectors harbor higher H3K9me3 and H3K27me3, otherwise, ATAC-seq, H3K4me3 and H3K36me3 is lowly on these type of genes.

(A). comparison of ATAC-seq and four histone methylation signal in core, secretome and RxLR genes. P values are calculated with the two-sample Kolmogorov-Smirnov test. (B), Representative snapshot of silenced pectinase and RxLR gene cluster. The snapshot was collected from IGV software, and specific loci were marked by blue frame.

S13 Fig. GO enrichment result of genes in OC, ST, H3K9DR and H3K27DR chromatin states.

