## Supplemental figures for "Genome-wide analyses of histone modifications and chromatin accessibility reveal the distinct genomic compartments in the Irish potato famine pathogen *Phytophthora infestans*"

#### S1 Fig.

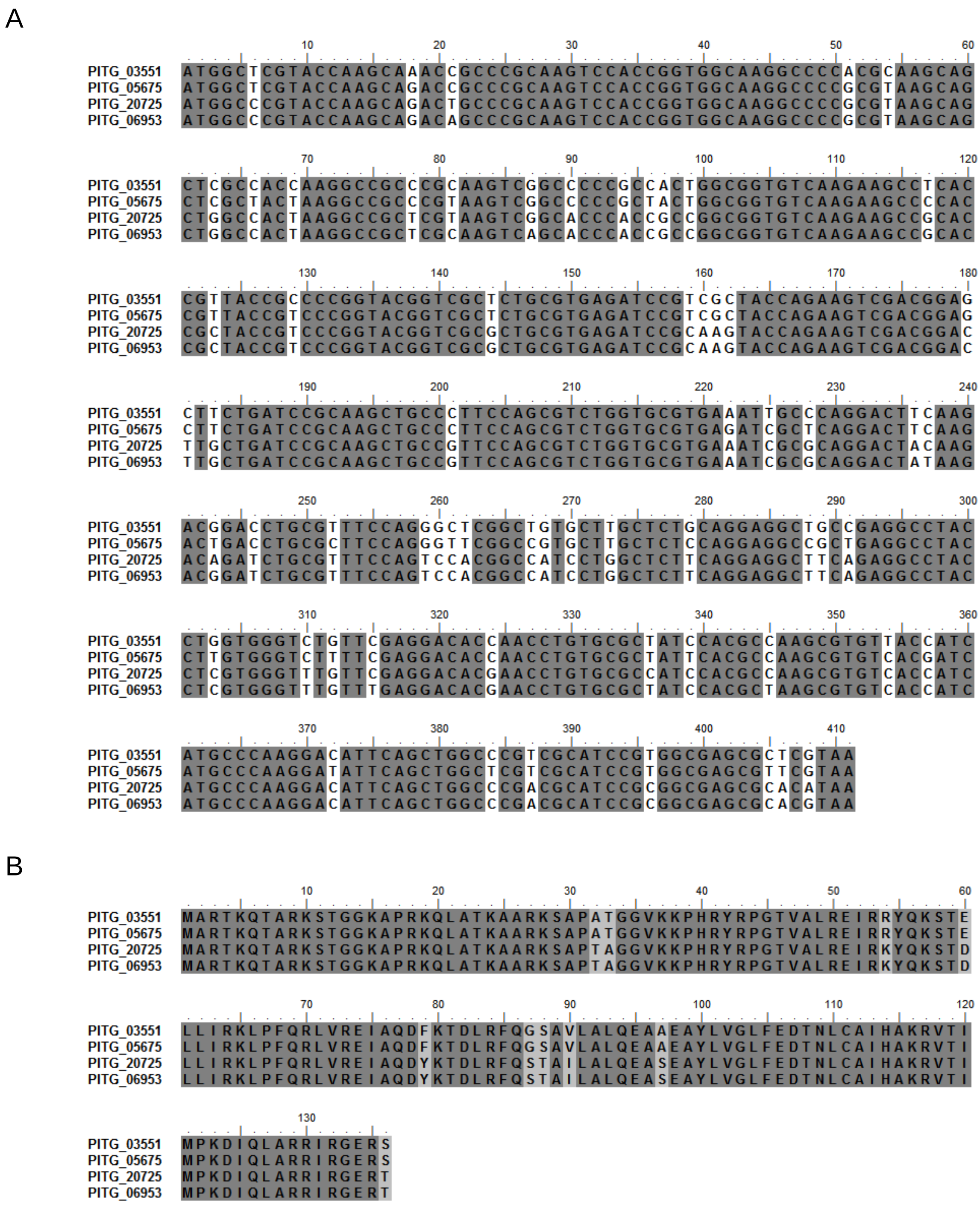
A
PITG_03551 ATGGCTCGTACCAAGCAAACCGCCCGCAAGTCCACCGGTGGCAAGGCCCCACGCAAGCAG
PITG_05675 ATGGCTCGTACCAAGCAGACCGCCCGCAAGTCCACCGGTGGCAAGGCCCCGCGTAAGCAG
PITG_20725 ATGGCCCGTACCAAGCAGACTGCCCGCAAGTCCACCGGTGGCAAGGCCCCGCGTAAGCAG
PITG_06953 ATGGCCCGTACCAAGCAGACAGCCCGCAAGTCCACCGGTGGCAAGGCCCCGCGTAAGCAG
PITG_03551 CTCGCCACCAAGGCCGCCCGCAAGTCGGCCCCCGCCACTGGCGGTGTCAAGAAGCCTCAC
PITG_05675 CTCGCTACTAAGGCCGCCCGTAAGTCGGCCCCCGCTACTGGCGGTGTCAAGAAGCCCCAC
PITG_20725 CTGGCCACTAAGGCCGCTCGTAAGTCGGCACCCACCGCCGGCGGTGTCAAGAAGCCGCAC
PITG_06953 CTGGCCACTAAGGCCGCTCGCAAGTCAGCACCCACCGCCGGCGGTGTCAAGAAGCCGCAC
PITG_03551 CGTTACCGCCCCGGTACGGTCGCTCTGCGTGAGATCCGTCGCTACCAGAAGTCGACGGAG
PITG_05675 CGTTACCGTCCCGGTACGGTCGCTCTGCGTGAGATCCGTCGCTACCAGAAGTCGACGGAG
PITG_20725 CGCTACCGTCCCGGTACGGTCGCGCTGCGTGAGATCCGCAAGTACCAGAAGTCGACGGAC
PITG_06953 CGCTACCGTCCCGGTACGGTCGCGCTGCGTGAGATCCGCAAGTACCAGAAGTCGACGGAC
PITG_03551 CTTCTGATCCGCAAGCTGCCCTTCCAGCGTCTGGTGCGTGAAATTGCCCAGGACTTCAAG
PITG_05675 CTTCTGATCCGCAAGCTGCCCTTCCAGCGTCTGGTGCGTGAGATCGCTCAGGACTTCAAG
PITG_20725 TTGCTGATCCGCAAGCTGCCGTTCCAGCGTCTGGTGCGTGAAATCGCGCAGGACTACAAG
PITG_06953 TTGCTGATCCGCAAGCTGCCGTTCCAGCGTCTGGTGCGTGAAATCGCGCAGGACTATAAG
PITG_03551 ACGGACCTGCGTTTCCAGGGCTCGGCTGTGCTTGCTCTGCAGGAGGCTGCCGAGGCCTAC
PITG_05675 ACTGACCTGCGCTTCCAGGGTTCGGCCGTGCTTGCTCTCCAGGAGGCCGCTGAGGCCTAC
PITG_20725 ACAGATCTGCGTTTCCAGTCCACGGCCATCCTGGCTCTTCAGGAGGCTTCAGAGGCCTAC
PITG_06953 ACGGATCTGCGTTTCCAGTCCACGGCCATCCTGGCTCTTCAGGAGGCTTCAGAGGCCTAC
PITG_03551 CTGGTGGGTCTGTTCGAGGACACCAACCTGTGCGCTATCCACGCCAAGCGTGTTACCATC
PITG_05675 CTTGTGGGTCTTTTCGAGGACACCAACCTGTGCGCTATTCACGCCAAGCGTGTCACGATC
PITG_20725 CTCGTGGGTTTGTTCGAGGACACGAACCTGTGCGCCATCCACGCCAAGCGTGTCACCATC
PITG_06953 CTCGTGGGTTTGTTTGAGGACACGAACCTGTGCGCTATCCACGCTAAGCGTGTCACCATC
PITG_03551 ATGCCCAAGGACATTCAGCTGGCCCGTCGCATCCGTGGCGAGCGCTCGTAA
PITG_05675 ATGCCCAAGGATATTCAGCTGGCTCGTCGCATCCGTGGCGAGCGTTCGTAA
PITG_20725 ATGCCCAAGGACATTCAGCTGGCCCGACGCATCCGCGGCGAGCGCACATAA
PITG_06953 ATGCCCAAGGACATTCAGCTGGCCCGACGCATCCGCGGCGAGCGCACGTAA
B
PITG_03551 MARTKQTARKSTGGKAPRKQLATKAARKSAPATGGVKKPHRYRPGTVALREIRRYQKSTE
PITG_05675 MARTKQTARKSTGGKAPRKQLATKAARKSAPATGGVKKPHRYRPGTVALREIRRYQKSTE
PITG_20725 MARTKQTARKSTGGKAPRKQLATKAARKSAPTAGGVKKPHRYRPGTVALREIRKYQKSTD
PITG_06953 MARTKQTARKSTGGKAPRKQLATKAARKSAPTAGGVKKPHRYRPGTVALREIRKYQKSTD
PITG_03551 LLIRKLPFQRLVREIAQDFKTDLRFQGSAVLALQEAAEAYLVGLFEDTNLCAIHAKRVTI
PITG_05675 LLIRKLPFQRLVREIAQDFKTDLRFQGSAVLALQEAAEAYLVGLFEDTNLCAIHAKRVTI
PITG_20725 LLIRKLPFQRLVREIAQDYKTDLRFQSTAILALQEASEAYLVGLFEDTNLCAIHAKRVTI
PITG_06953 LLIRKLPFQRLVREIAQDYKTDLRFQSTAILALQEASEAYLVGLFEDTNLCAIHAKRVTI
PITG_03551 MPKDIQLARRIRGERS
PITG_05675 MPKDIQLARRIRGERS
PITG_20725 MPKDIQLARRIRGERT
PITG_06953 MPKDIQLARRIRGERT

**S2 Fig.**

A

B

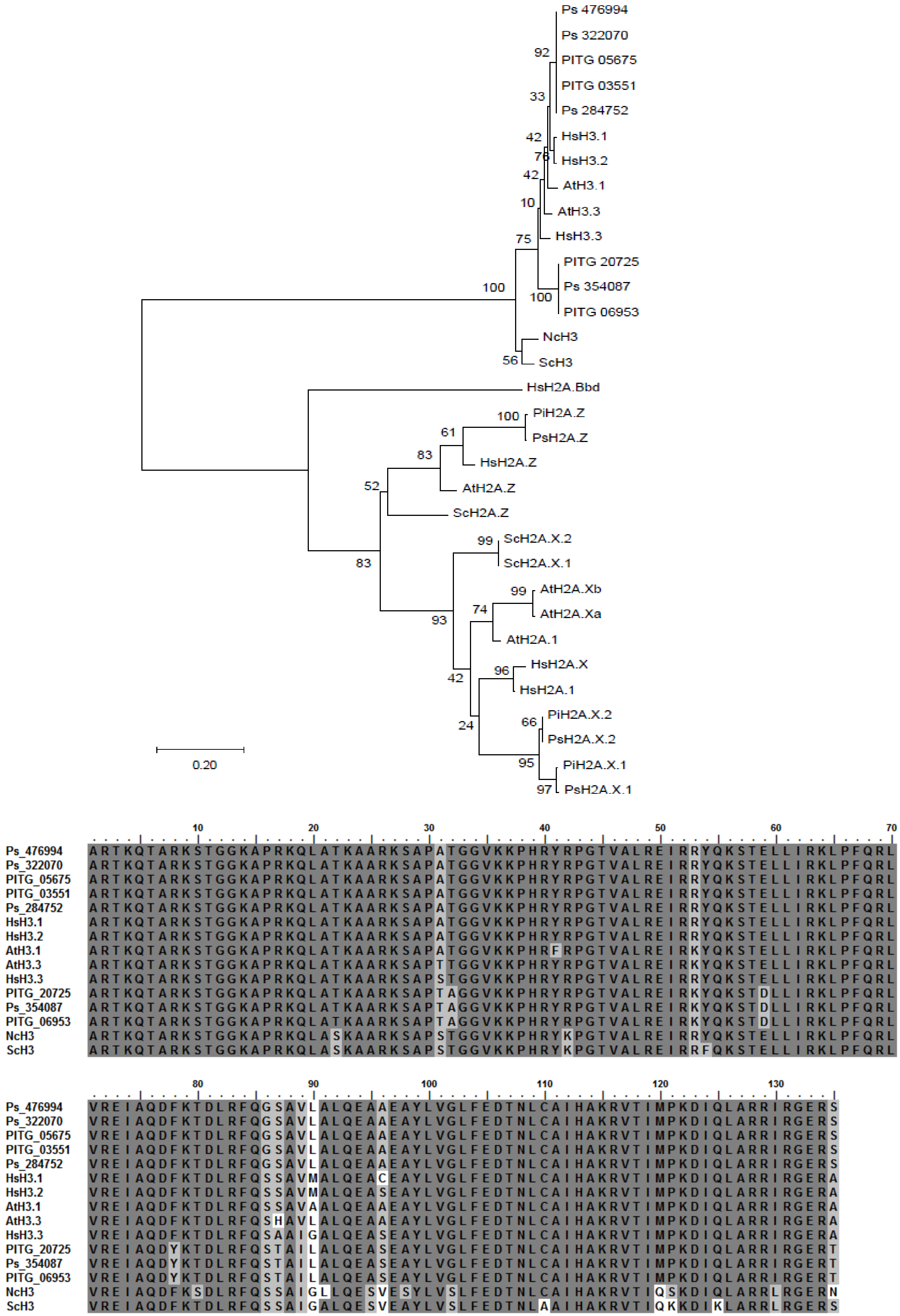
Ps 476994
Ps 322070
PITG 05675
PITG 03551
Ps 284752
HsH3.1
HsH3.2
AtH3.1
AtH3.3
HsH3.3
PITG 20725
Ps 354087
PITG 06953
NcH3
ScH3
HsH2A.Bbd
PiH2A.Z
PsH2A.Z
HsH2A.Z
AtH2A.Z
ScH2A.Z
ScH2A.X.2
ScH2A.X.1
AtH2A.Xb
AtH2A.Xa
AtH2A.1
HsH2A.X
HsH2A.1
PiH2A.X.2
PsH2A.X.2
PiH2A.X.1
PsH2A.X.1
92
33
42
76
42
10
75
100
100
56
100
61
83
52
83
99
99
74
93
96
42
24
66
95
97
0.20
10 20 30 40 50 60 70
Ps_476994 ARTKQTARKSTGGKAPRKQLATKAARKSAPATGGVKKPHRYRPGTVALREIRRYQKSTELLIRKLPFQRL
Ps_322070 ARTKQTARKSTGGKAPRKQLATKAARKSAPATGGVKKPHRYRPGTVALREIRRYQKSTELLIRKLPFQRL
PITG_05675 ARTKQTARKSTGGKAPRKQLATKAARKSAPATGGVKKPHRYRPGTVALREIRRYQKSTELLIRKLPFQRL
PITG_03551 ARTKQTARKSTGGKAPRKQLATKAARKSAPATGGVKKPHRYRPGTVALREIRRYQKSTELLIRKLPFQRL
Ps_284752 ARTKQTARKSTGGKAPRKQLATKAARKSAPATGGVKKPHRYRPGTVALREIRRYQKSTELLIRKLPFQRL
HsH3.1 ARTKQTARKSTGGKAPRKQLATKAARKSAPATGGVKKPHRYRPGTVALREIRRYQKSTELLIRKLPFQRL
HsH3.2 ARTKQTARKSTGGKAPRKQLATKAARKSAPATGGVKKPHRYRPGTVALREIRRYQKSTELLIRKLPFQRL
AtH3.1 ARTKQTARKSTGGKAPRKQLATKAARKSAPATGGVKKPHRFRPGTVALREIRRYQKSTELLIRKLPFQRL
AtH3.3 ARTKQTARKSTGGKAPRKQLATKAARKSAPTTGGVKKPHRYRPGTVALREIRKYQKSTELLIRKLPFQRL
HsH3.3 ARTKQTARKSTGGKAPRKQLATKAARKSAPSTGGVKKPHRYRPGTVALREIRRYQKSTELLIRKLPFQRL
PITG_20725 ARTKQTARKSTGGKAPRKQLATKAARKSAPTAGGVKKPHRYRPGTVALREIRKYQKSTDLLIRKLPFQRL
Ps_354087 ARTKQTARKSTGGKAPRKQLATKAARKSAPTAGGVKKPHRYRPGTVALREIRKYQKSTDLLIRKLPFQRL
PITG_06953 ARTKQTARKSTGGKAPRKQLATKAARKSAPTAGGVKKPHRYRPGTVALREIRKYQKSTDLLIRKLPFQRL
NcH3 ARTKQTARKSTGGKAPRKQLASKAARKSAPSTGGVKKPHRYKPGTVALREIRRYQKSTELLIRKLPFQRL
ScH3 ARTKQTARKSTGGKAPRKQLASKAARKSAPSTGGVKKPHRYKPGTVALREIRRFQKSTELLIRKLPFQRL
80 90 100 110 120 130
Ps_476994 VREIAQDFKTDLRFQGSAVLALQEAAEAYLVGLFEDTNLCAIHAKRVTIMPKDIQLARRIRGERS
Ps_322070 VREIAQDFKTDLRFQGSAVLALQEAAEAYLVGLFEDTNLCAIHAKRVTIMPKDIQLARRIRGERS
PITG_05675 VREIAQDFKTDLRFQGSAVLALQEAAEAYLVGLFEDTNLCAIHAKRVTIMPKDIQLARRIRGERS
PITG_03551 VREIAQDFKTDLRFQGSAVLALQEAAEAYLVGLFEDTNLCAIHAKRVTIMPKDIQLARRIRGERS
Ps_284752 VREIAQDFKTDLRFQGSAVLALQEAAEAYLVGLFEDTNLCAIHAKRVTIMPKDIQLARRIRGERS
HsH3.1 VREIAQDFKTDLRFQSSAVLALQEACEAYLVGLFEDTNLCAIHAKRVTIMPKDIQLARRIRGERA
HsH3.2 VREIAQDFKTDLRFQSSAVMALQEASEAYLVGLFEDTNLCAIHAKRVTIMPKDIQLARRIRGERA
AtH3.1 VREIAQDFKTDLRFQSSAVAALQEAAEAYLVGLFEDTNLCAIHAKRVTIMPKDIQLARRIRGERA
AtH3.3 VREIAQDFKTDLRFQSHAVLALQEAAEAYLVGLFEDTNLCAIHAKRVTIMPKDIQLARRIRGERA
HsH3.3 VREIAQDFKTDLRFQSAAIGALQEASEAYLVGLFEDTNLCAIHAKRVTIMPKDIQLARRIRGERA
PITG_20725 VREIAQDYKTDLRFQSTAILALQEASEAYLVGLFEDTNLCAIHAKRVTIMPKDIQLARRIRGERT
Ps_354087 VREIAQDYKTDLRFQSTAILALQEASEAYLVGLFEDTNLCAIHAKRVTIMPKDIQLARRIRGERT
PITG_06953 VREIAQDYKTDLRFQSTAILALQEASEAYLVGLFEDTNLCAIHAKRVTIMPKDIQLARRIRGERT
NcH3 VREIAQDFKSDLRFQSSAIGLLQESVESYLVSLFEDTNLCAIHAKRVTIQSKDIQLARRLRGERN
ScH3 VREIAQDFKTDLRFQSSAIGALQESVEAYLVSLFEDTNLAAIHAKRVTIQKKDIKLARRLRGERS

### S3 Fig.

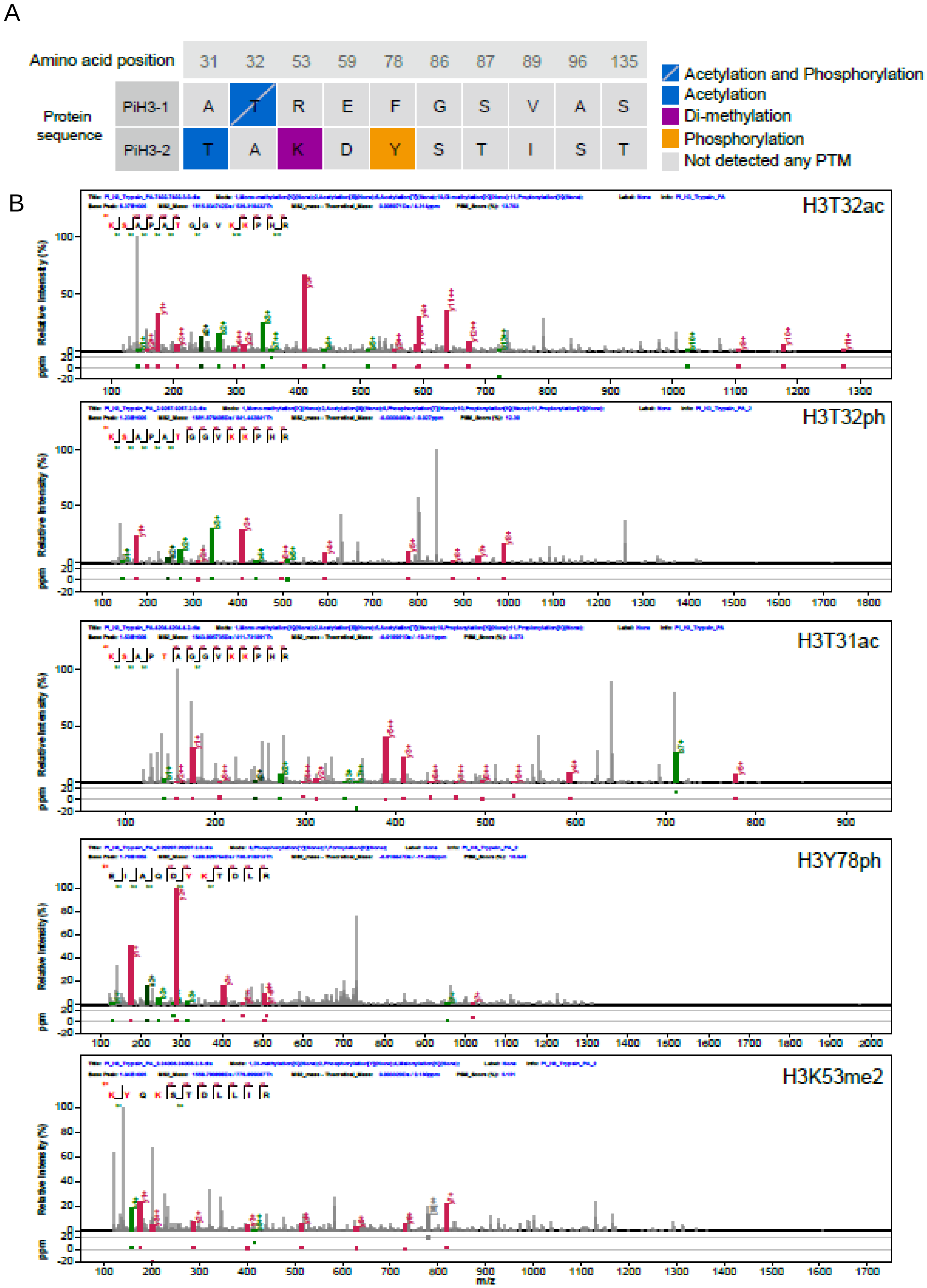
A
Amino acid position
31
32
53
59
78
86
87
89
96
135
Protein sequence
PiH3-1
A
T
R
E
F
G
S
V
A
S
PiH3-2
T
A
K
D
Y
S
T
I
S
T
Acetylation and Phosphorylation
Acetylation
Di-methylation
Phosphorylation
Not detected any PTM
B
H3T32ac
H3T32ph
H3T31ac
H3Y78ph
H3K53me2
Relative Intensity (%)
ppm
m/z

**S4 Fig.**

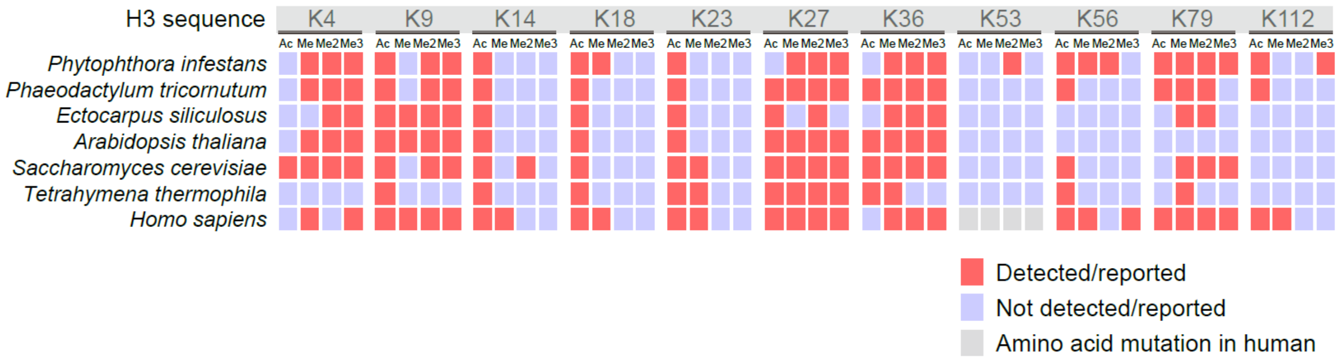
H3 sequence
K4
K9
K14
K18
K23
K27
K36
K53
K56
K79
K112
Ac Me Me2 Me3
Phytophthora infestans
Phaeodactylum tricornutum
Ectocarpus siliculosus
Arabidopsis thaliana
Saccharomyces cerevisiae
Tetrahymena thermophila
Homo sapiens
Detected/reported
Not detected/reported
Amino acid mutation in human

**S5 Fig.**

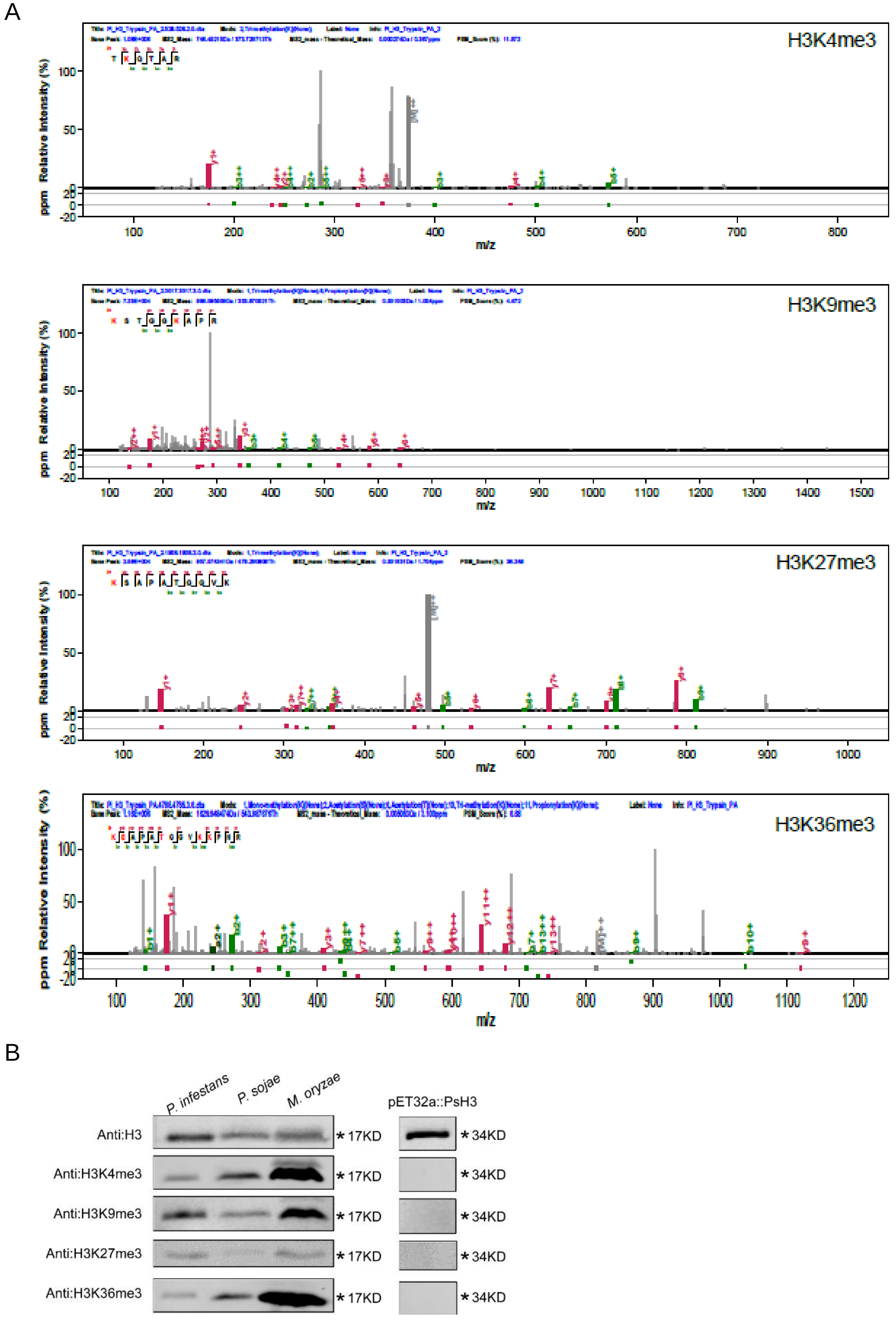
A
H3K4me3
H3K9me3
H3K27me3
H3K36me3
Relative Intensity (%)
ppm
m/z
B
P. infestans
P. sojae
M. oryzae
pET32a::PsH3
Anti:H3
Anti:H3K4me3
Anti:H3K9me3
Anti:H3K27me3
Anti:H3K36me3
17KD
34KD

**S6 Fig.**

A

| Sample | Total reads | Mapped reads | Mapping ratio |
| --- | --- | --- | --- |
| H3K4me3_rep1 | 33531634 | 32531877 | 97.02% |
| H3K4me3_rep2 | 36933784 | 35800560 | 96.93% |
| H3K36me3_rep1 | 42252867 | 41080249 | 97.22% |
| H3K36me3_rep2 | 21876722 | 21029788 | 96.13% |
| H3K9me3_rep1 | 37282908 | 36543538 | 98.02% |
| H3K9me3_rep2 | 36425754 | 35879658 | 98.50% |
| H3K27me3_rep1 | 21336100 | 20799956 | 97.49% |
| H3K27me3_rep2 | 19392173 | 18797262 | 96.93% |
| input_rep1 | 37787030 | 36811010 | 97.42% |
| input_rep2 | 21109505 | 20278096 | 96.06% |
| ATAC-seq_rep1 | 106657182 | 98051214 | 91.93% |
| ATAC-seq_rep2 | 107552620 | 98627857 | 91.70% |

B

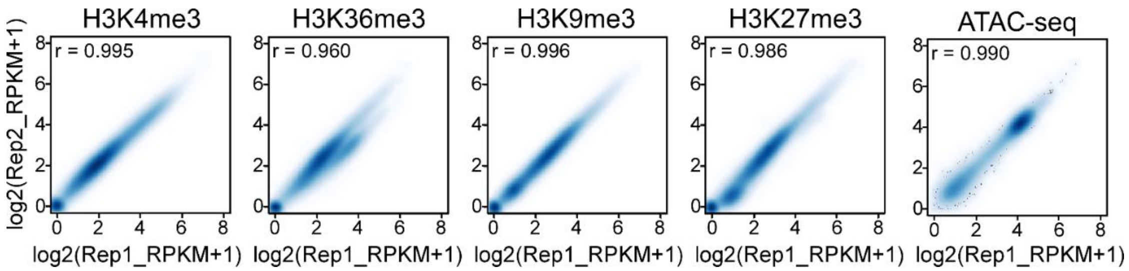


C

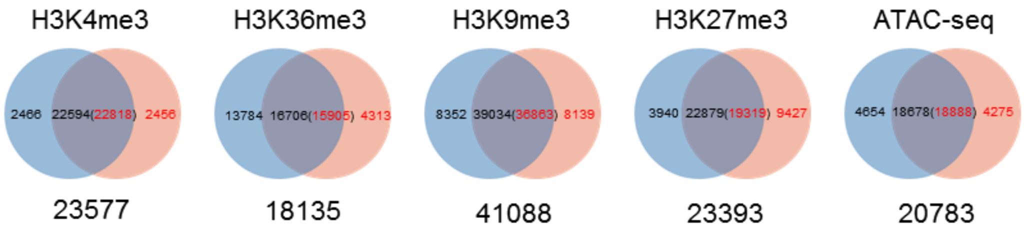

**S7 Fig.**

A

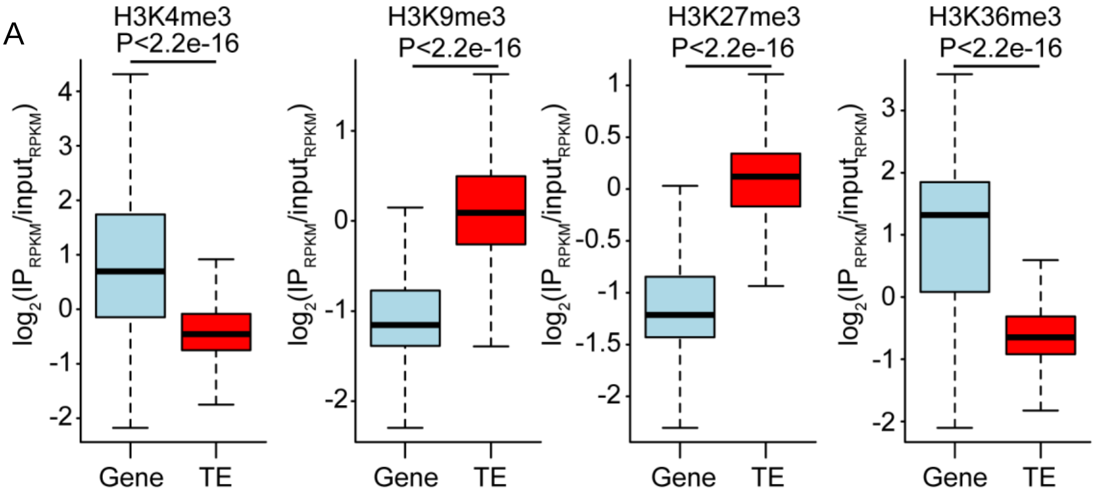


B

| | H3K9me3 | H3K27me3 | H3K4me3 | H3K36me3 | Expected |
|---|---|---|---|---|---|
| LTR | 0.55 | 0.66 | 0.11 | 0.04 | 0.30 |
| DNA transposon | 0.16 | 0.05 | 0.07 | 0.03 | 0.14 |
| non-LTR | 0.02 | 0.04 | 0.01 | 0.00 | 0.01 |

**S8 Fig.**

A

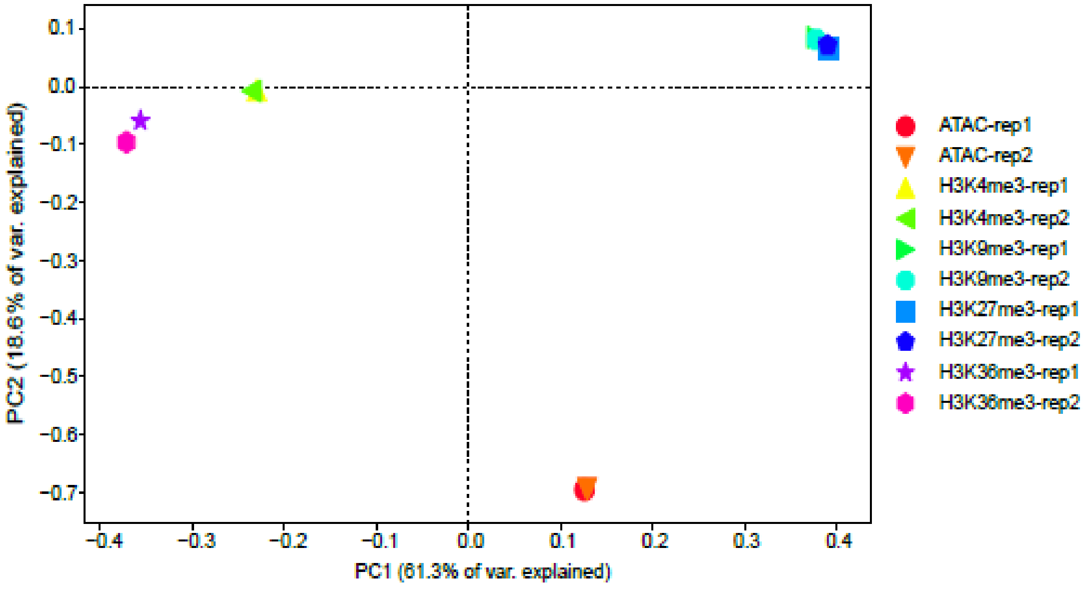


B

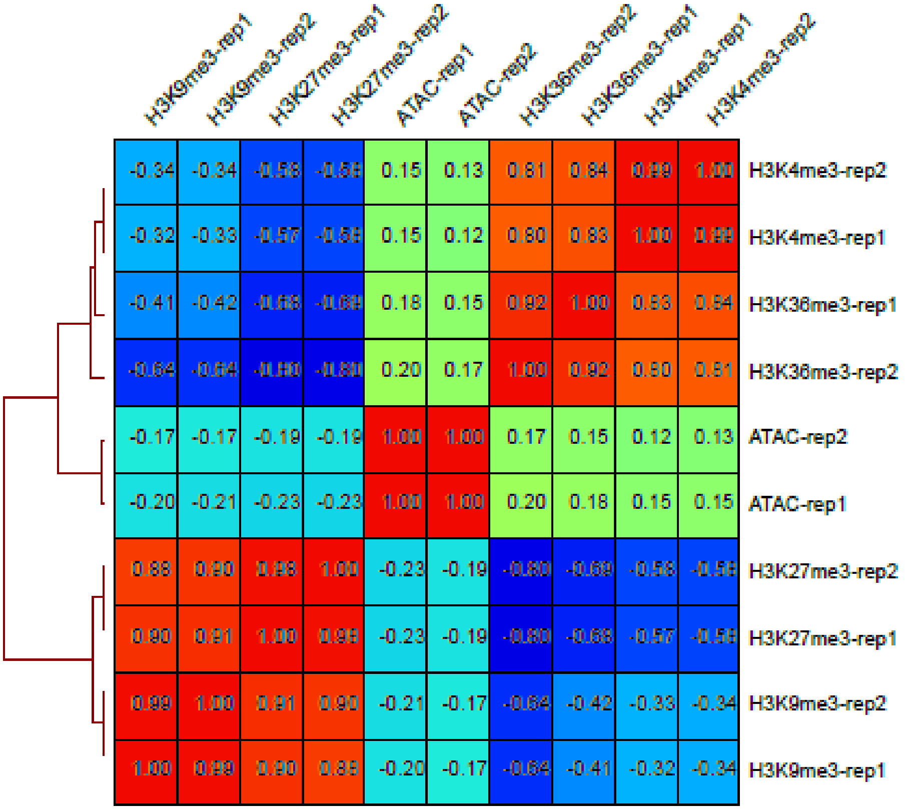


| | H3K9me3-rep1 | H3K9me3-rep2 | H3K27me3-rep1 | H3K27me3-rep2 | ATAC-rep1 | ATAC-rep2 | H3K36me3-rep2 | H3K36me3-rep1 | H3K4me3-rep1 | H3K4me3-rep2 |
|---|---|---|---|---|---|---|---|---|---|---|
| H3K4me3-rep2 | -0.34 | -0.34 | -0.58 | -0.58 | 0.15 | 0.13 | 0.81 | 0.84 | 0.99 | 1.00 |
| H3K4me3-rep1 | -0.32 | -0.33 | -0.57 | -0.58 | 0.15 | 0.12 | 0.80 | 0.83 | 1.00 | 0.99 |
| H3K36me3-rep1 | -0.41 | -0.42 | -0.68 | -0.69 | 0.18 | 0.15 | 0.92 | 1.00 | 0.83 | 0.84 |
| H3K36me3-rep2 | -0.64 | -0.64 | -0.80 | -0.80 | 0.20 | 0.17 | 1.00 | 0.92 | 0.80 | 0.81 |
| ATAC-rep2 | -0.17 | -0.17 | -0.19 | -0.19 | 1.00 | 1.00 | 0.17 | 0.15 | 0.12 | 0.13 |
| ATAC-rep1 | -0.20 | -0.21 | -0.23 | -0.23 | 1.00 | 1.00 | 0.20 | 0.18 | 0.15 | 0.15 |
| H3K27me3-rep2 | 0.88 | 0.90 | 0.98 | 1.00 | -0.23 | -0.19 | -0.80 | -0.69 | -0.58 | -0.58 |
| H3K27me3-rep1 | 0.90 | 0.91 | 1.00 | 0.98 | -0.23 | -0.19 | -0.80 | -0.68 | -0.57 | -0.58 |
| H3K9me3-rep2 | 0.99 | 1.00 | 0.91 | 0.90 | -0.21 | -0.17 | -0.64 | -0.42 | -0.33 | -0.34 |
| H3K9me3-rep1 | 1.00 | 0.99 | 0.90 | 0.88 | -0.20 | -0.17 | -0.64 | -0.41 | -0.32 | -0.34 |

**S9 Fig.**

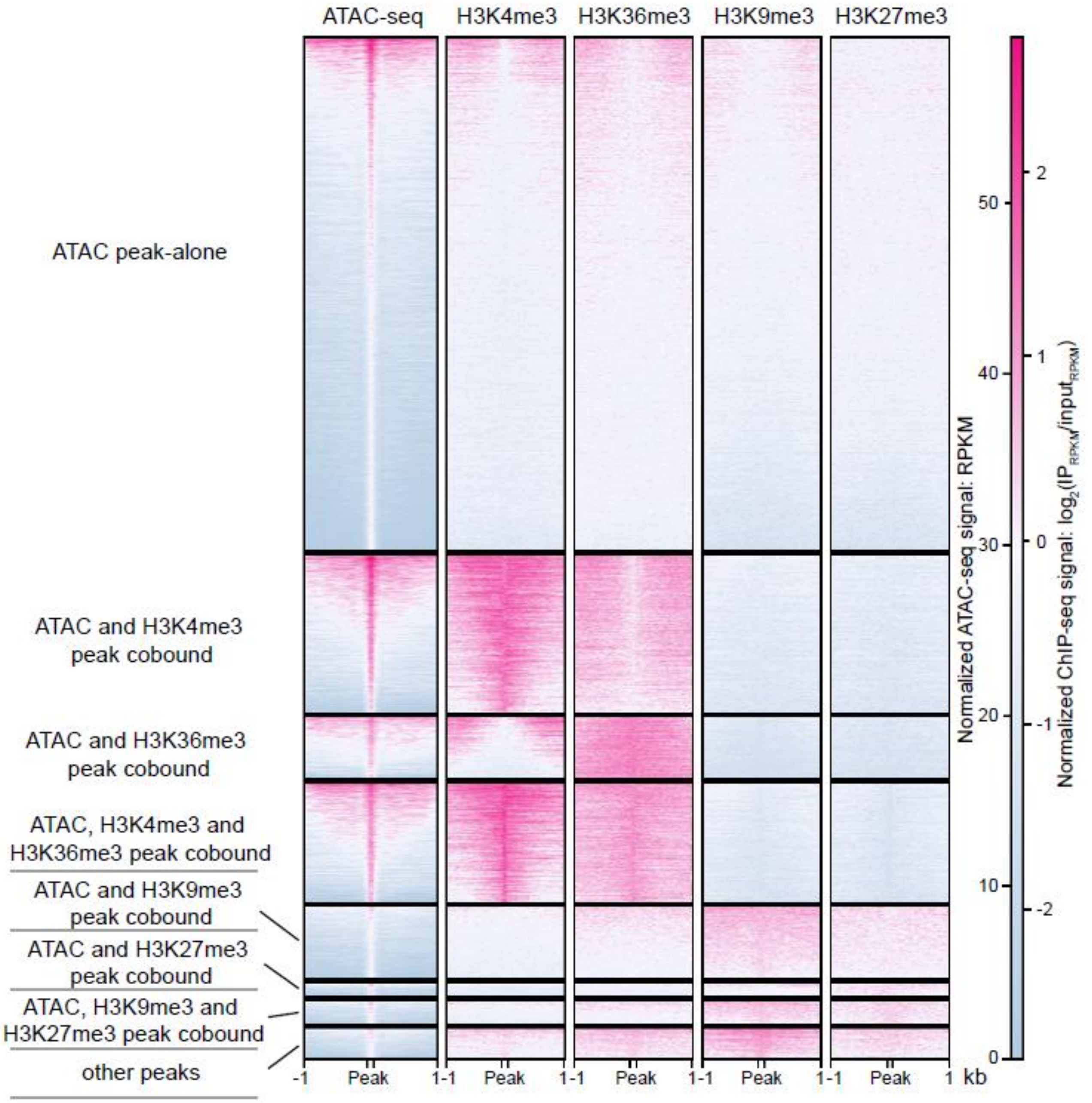

ATAC-seq
H3K4me3
H3K36me3
H3K9me3
H3K27me3
ATAC peak-alone
ATAC and H3K4me3 peak cobound
ATAC and H3K36me3 peak cobound
ATAC, H3K4me3 and H3K36me3 peak cobound
ATAC and H3K9me3 peak cobound
ATAC and H3K27me3 peak cobound
ATAC, H3K9me3 and H3K27me3 peak cobound
other peaks
-1
Peak
1
kb
Normalized ATAC-seq signal: RPKM
0
10
20
30
40
50
Normalized ChIP-seq signal: log2(IP_RPKM/input_RPKM)
-2
-1
0
1
2

**S10 Fig.**

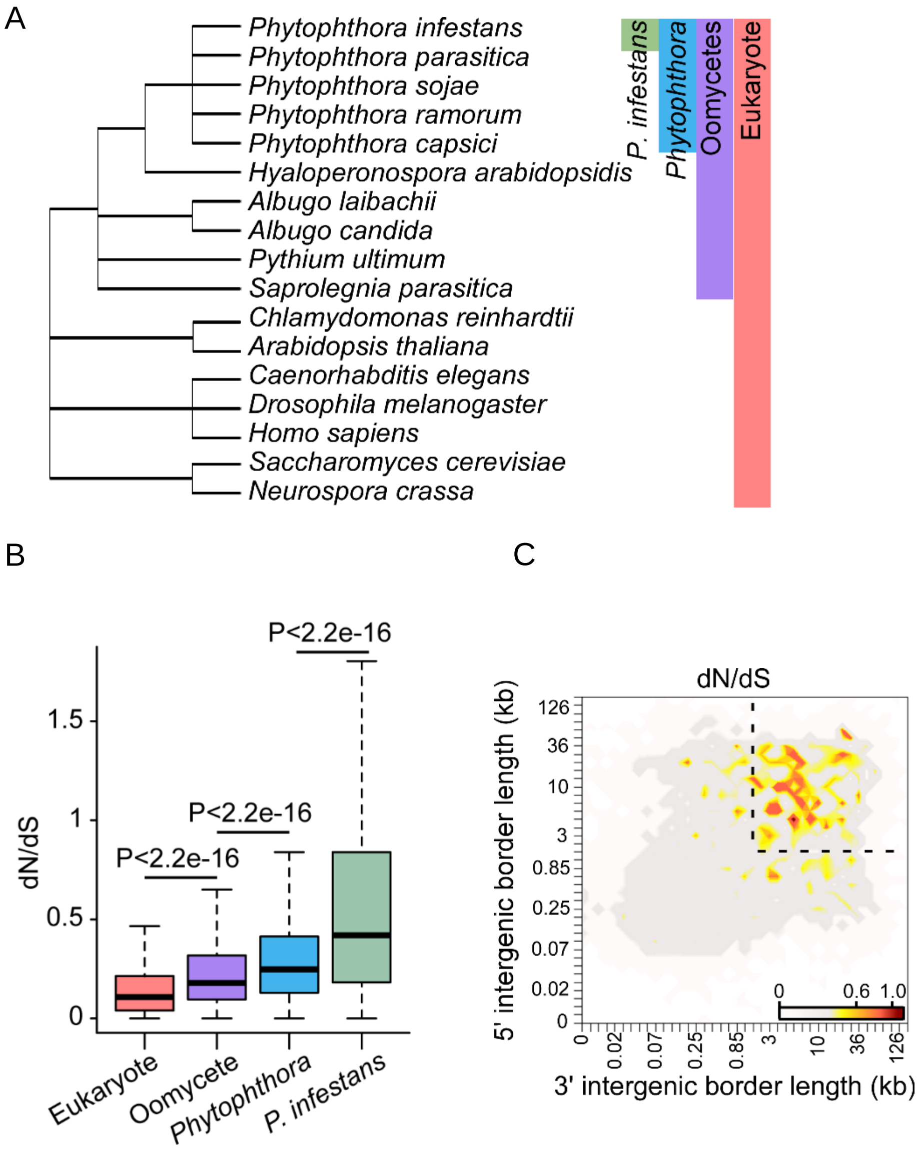
A
Phytophthora infestans
Phytophthora parasitica
Phytophthora sojae
Phytophthora ramorum
Phytophthora capsici
Hyaloperonospora arabidopsidis
Albugo laibachii
Albugo candida
Pythium ultimum
Saprolegnia parasitica
Chlamydomonas reinhardtii
Arabidopsis thaliana
Caenorhabditis elegans
Drosophila melanogaster
Homo sapiens
Saccharomyces cerevisiae
Neurospora crassa
P. infestans
Phytophthora
Oomycetes
Eukaryote
B
P<2.2e-16
P<2.2e-16
P<2.2e-16
dN/dS
1.5
1
0.5
0
Eukaryote
Oomycete
Phytophthora
P. infestans
C
dN/dS
5' intergenic border length (kb)
126
36
10
3
0.85
0.25
0.07
0.02
0
0.6
1.0
0.02
0.07
0.25
0.85
3
10
36
126
3' intergenic border length (kb)

**S11 Fig.**

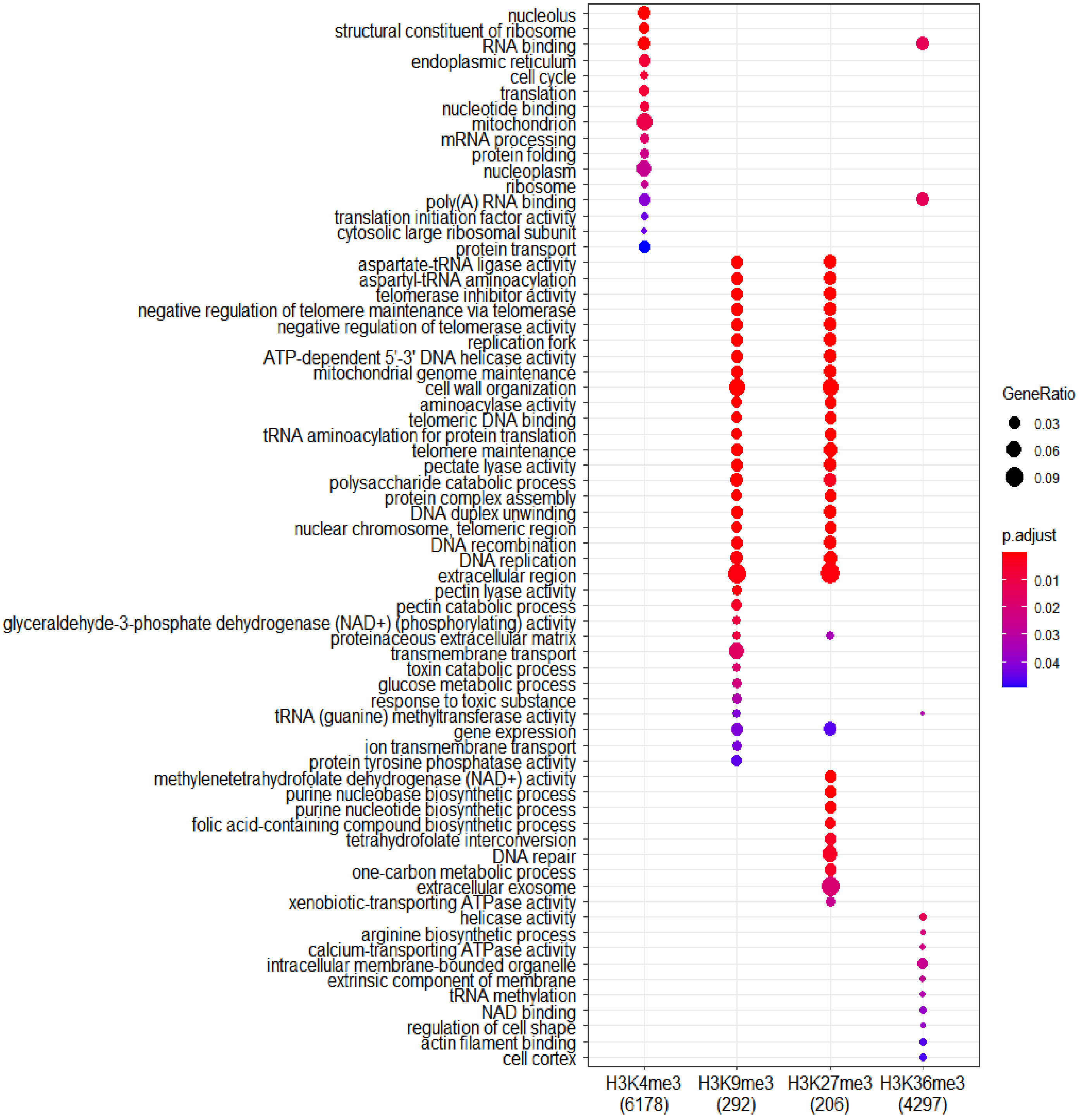
nucleolus
structural constituent of ribosome
RNA binding
endoplasmic reticulum
cell cycle
translation
nucleotide binding
mitochondrion
mRNA processing
protein folding
nucleoplasm
ribosome
poly(A) RNA binding
translation initiation factor activity
cytosolic large ribosomal subunit
protein transport
aspartate-tRNA ligase activity
aspartyl-tRNA aminoacylation
telomerase inhibitor activity
negative regulation of telomere maintenance via telomerase
negative regulation of telomerase activity
replication fork
ATP-dependent 5'-3' DNA helicase activity
mitochondrial genome maintenance
cell wall organization
aminoacylase activity
telomeric DNA binding
tRNA aminoacylation for protein translation
telomere maintenance
pectate lyase activity
polysaccharide catabolic process
protein complex assembly
DNA duplex unwinding
nuclear chromosome, telomeric region
DNA recombination
DNA replication
extracellular region
pectin lyase activity
pectin catabolic process
glyceraldehyde-3-phosphate dehydrogenase (NAD+) (phosphorylating) activity
proteinaceous extracellular matrix
transmembrane transport
toxin catabolic process
glucose metabolic process
response to toxic substance
tRNA (guanine) methyltransferase activity
gene expression
ion transmembrane transport
protein tyrosine phosphatase activity
methylenetetrahydrofolate dehydrogenase (NAD+) activity
purine nucleobase biosynthetic process
purine nucleotide biosynthetic process
folic acid-containing compound biosynthetic process
tetrahydrofolate interconversion
DNA repair
one-carbon metabolic process
extracellular exosome
xenobiotic-transporting ATPase activity
helicase activity
arginine biosynthetic process
calcium-transporting ATPase activity
intracellular membrane-bounded organelle
extrinsic component of membrane
tRNA methylation
NAD binding
regulation of cell shape
actin filament binding
cell cortex
H3K4me3 (6178)
H3K9me3 (292)
H3K27me3 (206)
H3K36me3 (4297)
GeneRatio
0.03
0.06
0.09
p.adjust
0.01
0.02
0.03
0.04

**S12 Fig.**

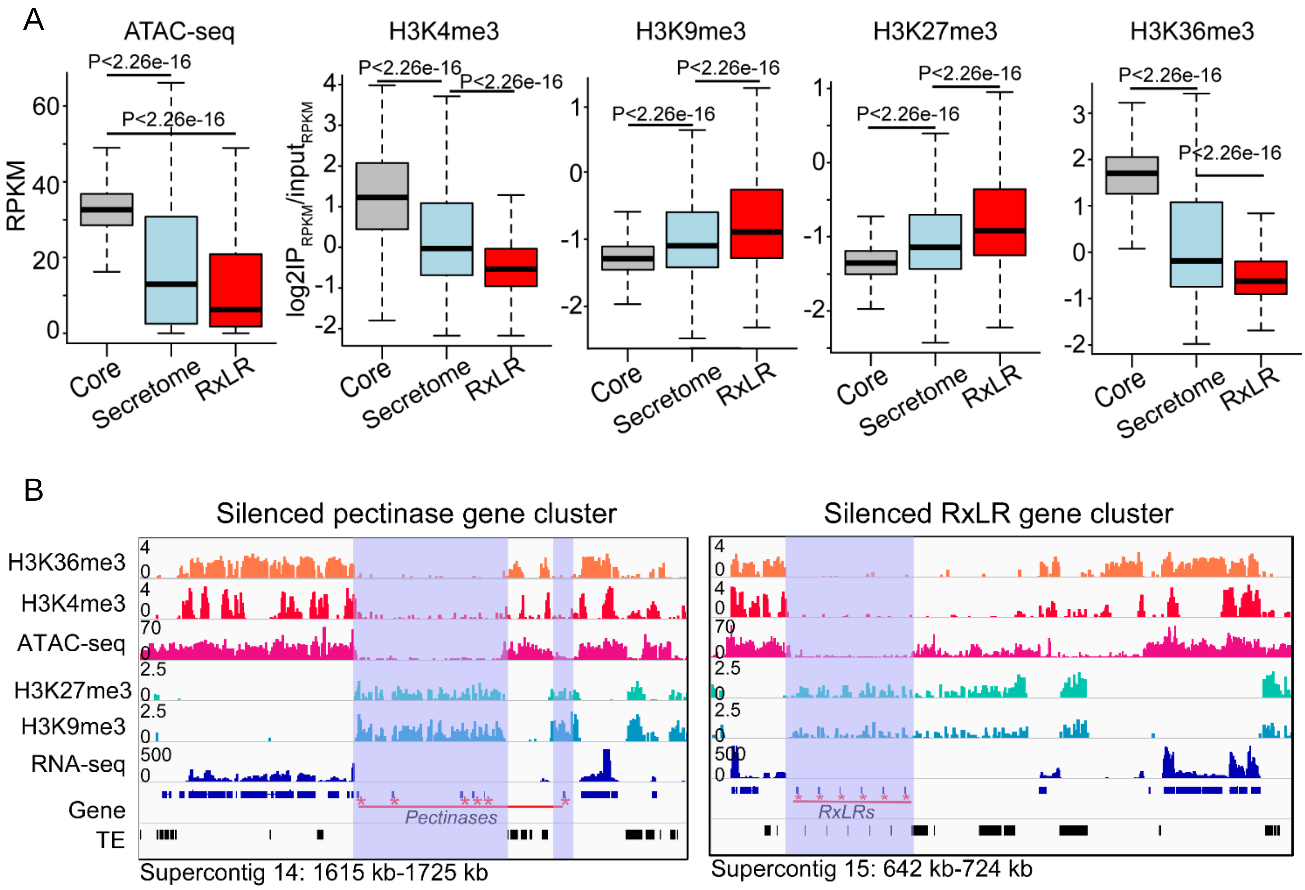
A
ATAC-seq
H3K4me3
H3K9me3
H3K27me3
H3K36me3
RPKM
log2IP_RPKM/input_RPKM
P<2.26e-16
Core
Secretome
RxLR
B
Silenced pectinase gene cluster
Silenced RxLR gene cluster
H3K36me3
H3K4me3
ATAC-seq
H3K27me3
H3K9me3
RNA-seq
Gene
TE
Pectinases
RxLRs
Supercontig 14: 1615 kb-1725 kb
Supercontig 15: 642 kb-724 kb

**S13 Fig.**

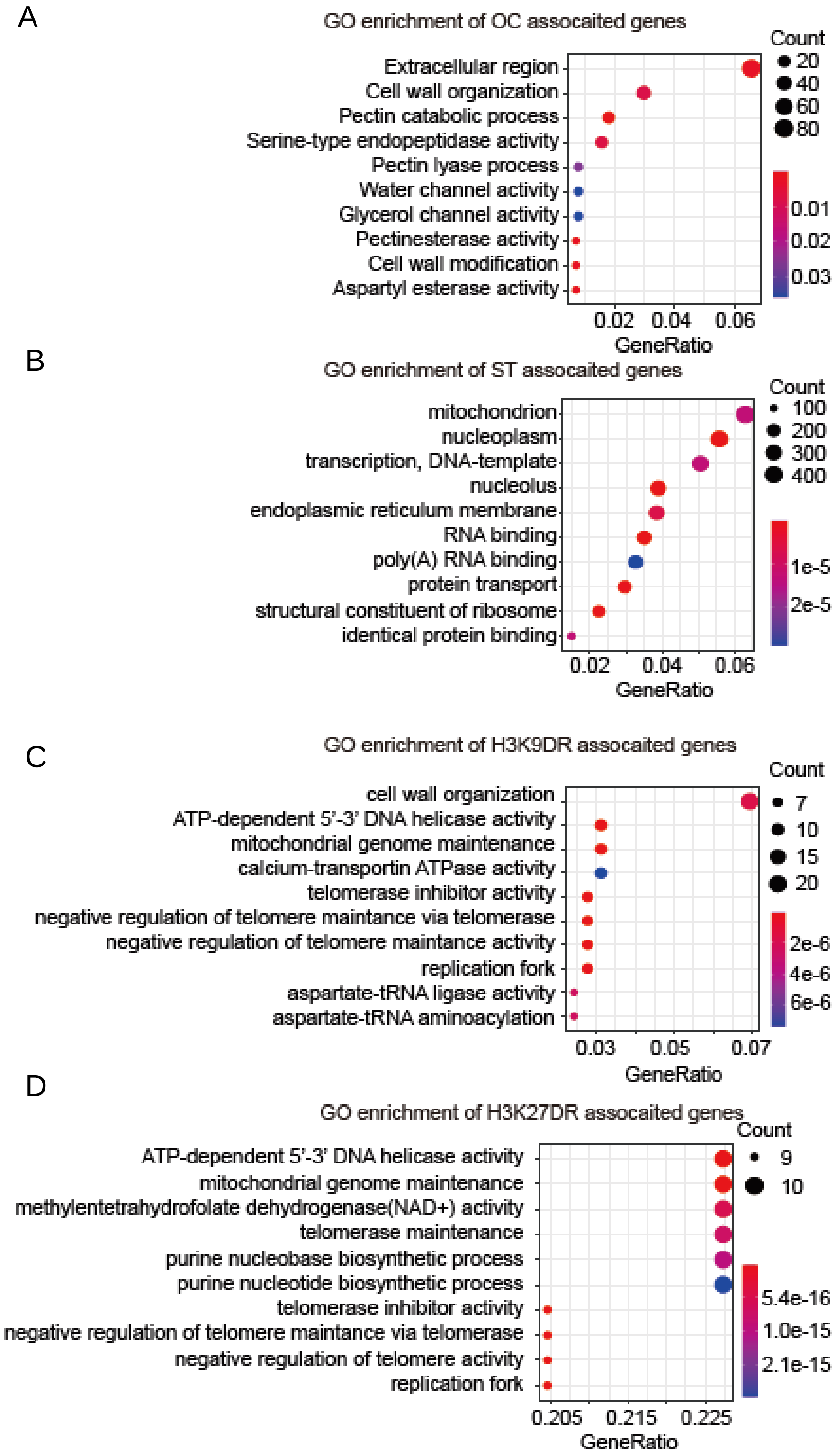
A
GO enrichment of OC assocaited genes
Extracellular region
Cell wall organization
Pectin catabolic process
Serine-type endopeptidase activity
Pectin lyase process
Water channel activity
Glycerol channel activity
Pectinesterase activity
Cell wall modification
Aspartyl esterase activity
0.02 0.04 0.06
GeneRatio
Count
20
40
60
80
0.01
0.02
0.03
B
GO enrichment of ST assocaited genes
mitochondrion
nucleoplasm
transcription, DNA-template
nucleolus
endoplasmic reticulum membrane
RNA binding
poly(A) RNA binding
protein transport
structural constituent of ribosome
identical protein binding
0.02 0.04 0.06
GeneRatio
Count
100
200
300
400
1e-5
2e-5
C
GO enrichment of H3K9DR assocaited genes
cell wall organization
ATP-dependent 5'-3' DNA helicase activity
mitochondrial genome maintenance
calcium-transportin ATPase activity
telomerase inhibitor activity
negative regulation of telomere maintance via telomerase
negative regulation of telomere maintance activity
replication fork
aspartate-tRNA ligase activity
aspartate-tRNA aminoacylation
0.03 0.05 0.07
GeneRatio
Count
7
10
15
20
2e-6
4e-6
6e-6
D
GO enrichment of H3K27DR assocaited genes
ATP-dependent 5'-3' DNA helicase activity
mitochondrial genome maintenance
methylentetrahydrofolate dehydrogenase(NAD+) activity
telomerase maintenance
purine nucleobase biosynthetic process
purine nucleotide biosynthetic process
telomerase inhibitor activity
negative regulation of telomere maintance via telomerase
negative regulation of telomere activity
replication fork
0.205 0.215 0.225
GeneRatio
Count
9
10
5.4e-16
1.0e-15
2.1e-15
